# Phenotypic and genomic characterization of *Pseudomonas aeruginosa* isolates recovered from catheter-associated urinary tract infections in an Egyptian hospital

**DOI:** 10.1101/2023.02.21.526938

**Authors:** Mohamed Eladawy, Jonathan C. Thomas, Lesley Hoyles

## Abstract

Catheter-associated urinary tract infections (CAUTIs) represent one of the major healthcare-associated infections, and *Pseudomonas aeruginosa* is a common Gram-negative bacterium associated with catheter infections in Egyptian clinical settings. The present study describes the phenotypic and genotypic characteristics of 31 *P. aeruginosa* isolates recovered from CAUTIs in an Egyptian hospital over a 3-month period. Genomes of isolates were of good quality and were confirmed to be *P. aeruginosa* by comparison to the type strain (average nucleotide identity, phylogenetic analysis). Clonal diversity among the isolates was determined; eight different sequence types were found (STs 244, 357, 381, 621, 773, 1430, 1667 and 3765), of which 357 and 773 are considered high-risk clones. Antimicrobial resistance (AMR) testing according to EUCAST guidelines showed the isolates were highly resistant to quinolones [ciprofloxacin (12/31, 38.7 %) and levofloxacin (9/31, 29 %) followed by tobramycin (10/31, 32.5 %)], and cephalosporins (7/31, 22.5 %). Genotypic analysis of resistance determinants predicted all isolates to encode a range of AMR genes, including those conferring resistance to aminoglycosides, β-lactamases, fluoroquinolones, fosfomycin, sulfonamides, tetracyclines and chloramphenicol. One isolate was found to carry a 422,938 bp pBT2436-like megaplasmid encoding OXA-520, the first report from Egypt of this emerging family of clinically important mobile genetic elements. All isolates were able to form biofilms, and were predicted to encode virulence genes associated with adherence, antimicrobial activity, antiphagocytosis, phospholipase enzymes, iron uptake, proteases, secretion systems, and toxins. The present study shows how phenotypic analysis alongside genomic analysis may help us understand the AMR and virulence profiles of *P. aeruginosa* contributing to CAUTIs in Egypt.

## INTRODUCTION

Urinary tract infections (UTIs) are among the most common bacterial infections that affect humans during their life span. They account for over 30 % of all health care-associated infections (HAIs) (Klevens et al., 2007). UTIs can be classified as uncomplicated or complicated depending on the site of infection and disease progress (Tan & Chlebicki, 2016). Urinary tract catheterization is a common practice which predisposes the host to complicated UTIs (Feneley et al., 2015). Instillation of a catheter in the urinary tract may cause mucosal-layer damage which disrupts the natural barrier and allows bacterial colonization (Kalsi et al., 2003).

*Pseudomonas aeruginosa* is an opportunistic pathogen that causes severe UTIs which are difficult to eradicate due to high intrinsic antimicrobial resistance (AMR) and the bacterium’s ability to develop new resistances during antibiotic treatment (Mittal et al., 2009). During the last decade there has been a dramatic worldwide increase in multidrug-resistant (MDR) *P. aeruginosa* responsible for various HAIs that lead to significant morbidity and mortality (Lamas Ferreiro et al., 2017). The World Health Organization named *P. aeruginosa* as a target of the highest priority for the development of new antibiotics (WHO, 2017). Infections caused by MDR *P. aeruginosa* were associated with a 70 % increase in cost per patient (Morales et al., 2012), and catheter-associated UTIs (CAUTI) cause an estimated 90,000 extra hospitalization days per year in the USA (Warren, 2001). According to the Centers for Disease Control and Prevention, more than 32,600 cases of HAIs were caused by MDR *P. aeruginosa* in the USA in 2017, which resulted in 2,700 deaths and $767M of estimated health-care costs (CDC, 2019).

In Egypt, mono-microbial infections represented 68.5 % of CAUTIs, while polymicrobial infections represented 31.43 % of catheterized patients admitted in 2021. Moreover, the prevalence of biofilm-dependent CAUTIs was about 82 %. The majority (81.25 %) of patients with catheters inserted for ≤14 days suffered from mono-bacterial colonization inside the catheter, and 42.11 % of patients with catheters inserted for one month had poly-microbial colonization (Ramadan et al., 2021).

There is an extensive variation in the epidemiology of MDR *P. aeruginosa* in the Middle East and North Africa (MENA) region in terms of AMR, prevalence, and genetic profiles. In general, there is high prevalence of MDR *P. aeruginosa* seen in Egypt (75.6 %) with similarities between neighboring countries, which might reflect comparable population and antibiotic-prescribing cultures (Al-Orphaly et al., 2021). However, there is no literature available on the genomic diversity of *P. aeruginosa* isolates contributing to CAUTIs in Egypt. We therefore aimed to investigate the genomic resistance and virulence profiles of *P. aeruginosa* contributing to CAUTIs by generating genome sequence data for isolates collected in an Egyptian hospital over a three-month period, and compared their genotypic and phenotypic data with respect to AMR profiles and biofilm-forming abilities.

## MATERIALS AND METHODS

### Recovery of isolates and ethical statement

Thirty-one *P. aeruginosa* isolates were recovered from urinary catheters (mono-microbial CAUTIs) between September and November 2021 by staff at the Urology and Nephrology Center, Mansoura, Egypt during routine diagnostic procedures (**Table (1)**). It was known that all isolates were associated with cases that had CAUTI as their primary diagnosis. We were informed that urine analysis had been performed on catheterized patients who presented with symptoms, mainly fever and dysuria. To collect a urine sample from patients with clinical signs/symptoms of a CAUTI, the urine had been aseptically aspirated from the urinary catheter and sent immediately to the hospital microbiology laboratory. Urine samples were examined under the microscope for white blood cells and processed using standard aseptic microbiological techniques. Urine samples were inoculated onto blood agar, Cystine-Lactose-Electrolyte-Deficient (CLED) agar, and MacConkey agar plates and incubated aerobically at 37 °C for up to three days. We were supplied with the isolates recovered on CLED agar, with only the date of isolation provided for samples in addition to confirmation of a CAUTI diagnosis; we were provided with no patient data.

**Table (1):** Summary information for the draft genomes generated from isolates described in this study.

| Isolate | Isolated | Genome accession | ANI (%) * | Coverage (x) # | Completeness (%) | Contamination (%) | Length (bp) | Contigs | GC content (%) | N50 | CDS | ST |
| --- | --- | --- | --- | --- | --- | --- | --- | --- | --- | --- | --- | --- |
| P1 | 23/9/2021 | JAPWLO000000000 | 99.28 | 104.7 | 99.68 | 0.14 | 7,090,567 | 33 | 65.8 | 670,701 | 6,518 | 244 |
| P2 | 23/9/2021 | JAPWLN000000000 | 99.21 | 54.1 | 99.68 | 0.24 | 7,561,602 | 176 | 65.4 | 218,286 | 6,967 | 244 |
| P3 | 23/9/2021 | JAPWLM000000000 | 99.25 | 91.8 | 99.68 | 0.14 | 7,089,819 | 33 | 65.8 | 671,466 | 6,519 | 244 |
| P4 | 27/9/2021 | JAPWLL000000000 | 99.25 | 49.8 | 99.68 | 0.11 | 6,567,076 | 29 | 66.3 | 731,473 | 5,990 | 381 |
| P5 | 27/9/2021 | JAPWLK000000000 | 98.71 | 42.8 | 99.68 | 0.15 | 6,872,195 | 141 | 66.0 | 281,948 | 6,358 | 773 |
| P6 | 27/9/2021 | JAPWLJ000000000 | 99.27 | 55.9 | 99.68 | 0.14 | 7,079,384 | 48 | 65.8 | 394,601 | 6,519 | 244 |
| P7 | 29/9/2021 | JAPWLI000000000 | 99.21 | 55.5 | 99.68 | 0.11 | 6,595,040 | 42 | 66.3 | 716,476 | 6,018 | 381 |
| P8 | 29/9/2021 | JAPWLH000000000 | 98.74 | 58.1 | 99.68 | 0.86 | 7,112,374 | 390 | 65.8 | 411,570 | 6,551 | 773 |
| P9 § | 5/10/2021 | JAPWLG000000000 | 99.29 | 56.0 | 100 | 0.02 | 6,990,601 | 3 | 65.8 | 6,518,599 | 6,556 | 3765 |
| P10 | 5/10/2021 | JAPWLF000000000 | 99.20 | 37.9 | 99.68 | 0.28 | 7,710,323 | 740 | 65.0 | 423,206 | 7,002 | 381 |
| P11 | 5/10/2021 | JAPWLE000000000 | 99.29 | 102.6 | 99.68 | 0.14 | 6,585,784 | 41 | 66.3 | 457,535 | 6,084 | 3765 |
| P12 | 11/10/2021 | JAPWLD000000000 | 99.24 | 41.3 | 99.68 | 0.11 | 6,589,324 | 58 | 66.3 | 656,238 | 6,011 | 381 |
| P13 | 11/10/2021 | JAPWLC000000000 | 99.26 | 69.7 | 99.68 | 0.16 | 6,492,143 | 42 | 66.2 | 427,633 | 5,924 | 1667 |
| P14 | 11/10/2021 | JAPWLB000000000 | 98.73 | 67.5 | 99.68 | 0.21 | 6,844,752 | 71 | 66.1 | 433,376 | 6,327 | 773 |
| P15 | 16/10/2021 | JAPWLA000000000 | 99.29 | 68.4 | 99.68 | 0.14 | 6,577,280 | 114 | 66.3 | 456,538 | 6,009 | 3765 |
| P16 | 16/10/2021 | JAPWKZ000000000 | 98.78 | 46.7 | 99.68 | 0.55 | 7,019,039 | 286 | 65.7 | 369,447 | 6,394 | 357 |
| P17 | 17/10/2021 | JAPWKY000000000 | 99.14 | 57.2 | 99.68 | 0.60 | 6,845,094 | 62 | 65.9 | 327,266 | 6,241 | 621 |
| P18 | 17/10/2021 | JAPWKX000000000 | 99.24 | 57.5 | 99.68 | 0.11 | 6,577,155 | 29 | 66.3 | 810,963 | 5,993 | 381 |
| P19 § | 20/10/2021 | JAPWKW000000000 | 99.22 | 43.7 | 100 | 0.18 | 6,632,993 | 3 | 66.2 | 5,895,732 | 6,036 | 381 |
| P20 | 20/10/2021 | JAPWKV000000000 | 98.72 | 66.1 | 99.68 | 0.21 | 6,835,420 | 70 | 66.1 | 316,419 | 6,320 | 773 |
| P22 | 20/10/2021 | JAPWKU000000000 | 99.28 | 74.9 | 99.68 | 0.14 | 7,082,297 | 36 | 65.8 | 670,701 | 6,522 | 244 |
| P23 § | 26/10/2021 | JAPWKT000000000 | 99.14 | 36.7 | 100 | 0.68 | 6,931,140 | 1 | 65.8 | 6,931,140 | 6,287 | 621 |
| P24 § | 26/10/2021 | JAPWKS000000000 | 99.23 | 43.2 | 100 | 0.18 | 6,688,005 | 6 | 66.2 | 5,887,181 | 6,109 | 381 |
| P25 | 26/10/2021 | JAPWKR000000000 | 98.78 | 83.2 | 99.68 | 0.50 | 6,642,761 | 33 | 66.2 | 457,730 | 6,034 | 357 |
| P26 | 26/10/2021 | JAPWKQ000000000 | 98.72 | 46.5 | 99.68 | 0.21 | 6,827,640 | 99 | 66.1 | 307,141 | 6,306 | 773 |
| P27 | 27/10/2021 | JAPWKP000000000 | 98.69 | 45.5 | 99.68 | 0.66 | 7,152,409 | 161 | 65.7 | 271,243 | 6,601 | 773 |
| P28 | 27/10/2021 | JAPWKO000000000 | 99.36 | 40.7 | 99.68 | 0.22 | 6,410,783 | 55 | 66.4 | 322,863 | 5,852 | 1430 |
| P29 | 1/11/2021 | JAPWKN000000000 | 99.35 | 42.4 | 99.68 | 0.17 | 6,757,213 | 145 | 66.1 | 400,482 | 6,200 | 3765 |
| P30 | 1/11/2021 | JAPWKM000000000 | 98.74 | 48.9 | 99.68 | 0.21 | 6,836,605 | 79 | 66.1 | 411,378 | 6,322 | 773 |
| P31 | 1/11/2021 | JAPWKL000000000 | 98.70 | 66.2 | 99.68 | 0.60 | 7,132,296 | 192 | 65.7 | 301,000 | 6,570 | 357 |
| P32 | 1/11/2021 | JAPWKK000000000 | 98.72 | 54.9 | 99.68 | 0.50 | 6,665,983 | 94 | 66.1 | 383,436 | 6,057 | 357 |

\* Illumina-only assemblies compared (fastANI) with the genome of *P. aeruginosa* DSM 50071<sup>T</sup>, NCBI Genome Assembly
GCF\_012987025.

The study of anonymized clinical isolates beyond the diagnostic requirement was approved by the Urology and Nephrology Center, Mansoura, Egypt. No other ethical approval was required for the use of the clinical isolates.

### Antimicrobial susceptibility testing

Antimicrobial susceptibility testing was performed using the disc diffusion test (DDT) on Mueller-Hinton agar (Oxoid Ltd, UK), with overnight cultures diluted to be equal to 0.5 McFarland standard (OD_600_ = 0.08–0.13) and spread (swabs) on the plates, followed by incubation at 37 °C for 18 h. Inhibition zone diameters were determined and recorded according to breakpoints of EUCAST guidelines 2022. *P. aeruginosa* ATCC 27853 was used as the standard strain.

### Assay of biofilm formation

The assay was performed as described previously (Eladawy et al., 2021; Merritt et al., 2005; Stepanovic et al., 2000). In brief, a single colony of each isolate was inoculated in 5 ml of tryptone soy broth (Oxoid Ltd) supplemented with 1 % glucose (TSBG). Cultures were incubated aerobically for 24 h at 37 °C without shaking. The overnight cultures were diluted to 1:100 using TSBG, then aliquots (100 µl) of the diluted cultures were introduced into wells of a 96-well plate. The plates were incubated aerobically for 24 h at 37 °C without shaking. Then, the spent medium was carefully removed from each well. The wells were washed three times with 200 µl sterile phosphate-buffered saline (pH 7.4; Oxoid Ltd) to remove any non-adherent planktonic cells. The adherent cells were fixed by heat treatment at 60 °C for 60 min to prevent widespread detachment of biofilms prior to dye staining. The adhered biofilms were then stained by addition of 1 % (w/v) crystal violet (150 µl per well) and the 96-well plate was left to incubate for 20 min. The excess stain was then carefully removed from the wells and discarded. The 96-well plate was carefully rinsed with distilled water three times, then the plate was inverted and left at room temperature until the wells were dry. The stained biofilms were solubilized by adding 33 % (v/v) glacial acetic acid (Sigma Aldrich) to each well (150 µl per well). After solubilization of stained biofilms, the OD_540_ was measured and recorded for all samples using a BioTek Cytation imaging reader spectrophotometer.

Un-inoculated medium was used as a negative control in biofilm assays. Biological (*n*=3) and technical (*n*=4) replicates were done for all isolates. *Salmonella enterica* serovar Enteritidis 27655S was used as a negative control in biofilm assays (Hayward et al., 2016).

### DNA extraction and whole-genome sequencing

For each isolate, a 500 µl aliquot of an overnight culture grown in nutrient broth (Oxoid Ltd) was used for DNA extraction using the Gentra Puregene Yeast/Bact. Kit (Qiagen) according to the manufacturer’s instructions. Quality and quantity of the extracted DNA were checked by NanoDrop™ 2000/2000c (ThermoFisher Scientific).

Illumina sequencing (Nextera XT Library Prep Kit; HiSeq/NovaSeq; 2 ×250 bp pairedend reads; min. 30× coverage) was performed by microbesNG (Birmingham, United Kingdom). Reads were adapter-trimmed to a minimum length of 36 nt using Trimmomatic 0.30 (Bolger et al., 2014) with a sliding window quality cut-off of Q15. *De novo*-assembled genomes (SPAdes v3.7; (Bankevich et al., 2012)) were returned to us by microbesNG.

Genomic DNA for four isolates (P9, P19, P23 and P24) was further sequenced to obtain long-read sequences using an Oxford Nanopore Technologies (ONT) MinION. The ligation sequencing kit SQK-LSK109 and native barcoding kit EXP-NBD104 were used for Nanopore library preparation. Libraries were loaded onto a MinION R9.4.1 flow cell and run for 48 h.

Fast5 files were basecalled using the SUP (super high accuracy) model of Guppy v6.4.2 and subsequently demultiplexed. Porechop (https://github.com/rrwick/Porechop) was used to trim end and middle adapter sequences and reads shorter than 1 kbp were discarded using Filtlong v0.2.1 (https://github.com/rrwick/Filtlong). Nanopore reads were *de novo* assembled using Flye v2.9.1 (Kolmogorov et al., 2019). Closed genomes were manually reoriented to begin with *dnaA*, prior to polishing with both Nanopore and Illumina reads. Assembled sequences were polished with Nanopore reads using four iterations of Racon v1.5.0 (Vaser et al., 2017), followed by Medaka v1.7.2 and Homopolish v0.3.4 (Huang et al., 2021). Resulting sequences were then polished with Illumina reads using Polypolish v0.5.0 (Wick & Holt, 2022), POLCA from the MaSuRCA v4.0.9 package (Zimin & Salzberg, 2020) and Nextpolish v1.4.1 (Hu et al., 2019).

### Bioinformatic analyses

Contigs with fewer than 500 bp were filtered from draft genomes using reformat.sh of BBmap 38.97 (Bushnell, 2014). CheckM v1.2.1 was used to assess genome quality (Parks et al., 2015). Identity of isolates as *P. aeruginosa* was confirmed by average nucleotide identity analysis (ANI) (fastANI v1.3.3) (Jain et al., 2018b) against the genome of the type strain of the species (DSM 50071^T^, NCBI Genome Assembly GCF_012987025.1). Bakta v1.5.1 (database v4.0) was used for annotating genes within genomes (Schwengers et al., 2021). The Bakta-annotated whole-genome sequence data are available from figshare in GenBank format. The Virulence Factor Database (Chen et al., 2005) was used to predict virulence genes encoded within genomes. Multilocus sequence type (MLST) of each isolate was determined using the MLST schema for *P. aeruginosa* at PubMLST (http://pubmlst.org/paeruginosa) (Curran et al., 2004; Jolley et al., 2018). PubMLST summary data were downloaded for 8,435 isolates on 16 December 2022. Antimicrobial resistance markers were identified using Resistance Gene Identifier (RGI) v6.0.0 tool of the Comprehensive Antibiotic Resistance Database (CARD) v3.2.5 (McArthur et al., 2013). Only resistance genes that showed a perfect or strict match with coverage for a given gene in the database are reported in this study. Phylogenetic analysis of genomic data was carried out using PhyloPhlAn 3.0 (Asnicar et al., 2020) with 245 *Pseudomonas* reference sequences downloaded from the Genome Taxonomy Database, release 07-RS207 (Supplementary Material: gtdb-search.csv) (Parks et al., 2018).

A BLASTN search was made using the megaplasmid pBT2436-like core gene sequences (*repA, parA, virB4*) described by (Cazares et al., 2020) against the contigs of our newly generated short-read genome sequence data. In addition, the reads from our short-read sequence data were trimmed to ≥70 nt each using cutadapt v4.1 (Martin, 2011) then mapped using BWA-MEM v.0.7.17-r1188 (Li, 2013) against the reference megaplasmid sequences shown in **Table (2)**. The presence of pBT2436-like megaplasmids in our genomes was assessed based on the percentage of reads mapped to the reference genomes of (Cazares et al., 2020) as extracted from the alignment files with samtools v.1.16.1 (Li et al., 2009). plaSquid was used to further characterize the plasmids (Giménez et al., 2022).

**Table (2):** pBT2436-like megaplasmid reference sequences included in this study.

| Plasmid | Species and strain | Size (bp) | No. of predicted genes * | Country | Source | GenBank accession | Reference(s) |
| --- | --- | --- | --- | --- | --- | --- | --- |
| pBT2436 | *P. aeruginosa* 2436 | 422,811 | 537 | Thailand | Respiratory infection | CP039989 | (Cazares et al., 2020) |
| pBT2101 | *P. aeruginosa* 2101 | 439,744 | 556 | Thailand | Respiratory infection | CP039991 | (Cazares et al., 2020) |
| unnamed2 | *P. aeruginosa* AR_0356 | 438,531 | 557 | Unknown | Unknown | CP027170 | (Cazares et al., 2020) |
| unnamed2 | *P. aeruginosa* AR439 | 437,392 | 549 | Unknown | Unknown | CP029096 | (Cazares et al., 2020) |
| unnamed3 | *P. aeruginosa* AR441 | 438,529 | 560 | Unknown | Unknown | CP029094 | (Cazares et al., 2020) |
| pJB37 | *P. aeruginosa* FFUP_PS_37 | 464,804 | 597 | Portugal | Respiratory infection | KY494864 | (Botelho et al., 2017; Cazares et al., 2020) |
| pBM413 | *P. aeruginosa* PA121617 | 423,017 | 537 | China | Respiratory infection | CP016215 | (Cazares et al., 2020; M. Li et al., 2022) |
| pOZ176 | *P. aeruginosa* PA96 | 500,839 | 621 | China | Respiratory infection | KC543497 | (Cazares et al., 2020; Xiong et al., 2013) |
| p12939-OXA | *P. aeruginosa* (unknown) | 496,436 | 607 | China | Unknown | MF344569 | (Cazares et al., 2020) |
| p727-IMP | *P. aeruginosa* (unknown) | 430,173 | 534 | China | Unknown | MF344568 | (Cazares et al., 2020) |
| pA681-IMP | *P. aeruginosa* (unknown) | 397,519 | 486 | China | Unknown | MF344570 | (Cazares et al., 2020) |
| pR31014-IMP | *P. aeruginosa* (unknown) | 374,000 | 456 | China | Unknown | MF344571 | (Cazares et al., 2020) |
| pRBL16 | *P. citronellolis* SJTE-3 | 370,338 | 486 | China | Wastewater sludge | CP015879 | (Cazares et al., 2020; Zheng et al., 2016) |
| p1 | *P. koreensis* P19E3 | 467,568 | 598 | Switzerland | *Origanum majorana* | CP027478 | (Cazares et al., 2020; Schmid et al., 2018) |
| pSY153-MDR | *P. putida* SY153 | 468,170 | 579 | China | Urinary tract infection | KY883660 | (Cazares et al., 2020; Yuan et al., 2017) |

\* Predicted in this study using Bakta.

Complete *Pseudomonas* plasmid sequences were downloaded from NCBI Genome on 19 December 2022 (Supplementary Material: plasmids.csv), and filtered to retain genomes >200,000 bp. These sequences were subject to BLASTN searches against the pBT2436 sequences for *repA, parA, virB4* as described above. Those plasmid sequences returning singlecopy hits for the three genes were subject to further analyses as follows.

For comparative analyses, the megaplasmid sequences were annotated using Bakta as described above for the *Pseudomonas* genome sequences. The Bakta-annotated plasmid sequence data are available from figshare in GenBank format. FastANI v1.33 (Jain et al., 2018a) was used to determine how similar the sequences of the newly identified megaplasmids were to those of pBT2436 and the other reference genomes (**Table (2)**); visualization of the conserved regions between pairs of plasmid sequences was achieved using the --visualize option of FastANI and the R script available at https://github.com/ParBLiSS/FastANI. The protein sequences predicted to be encoded by all the plasmids were concatenated, sorted by length (longest to shortest) using vsearch v2.15.2_linux_x86_64 (Rognes et al., 2016) and clustered using MMseqs2 v13.45111 (Steinegger & Söding, 2017) (80 % identity, 80 % coverage). Those core sequences found in MMseqs2 clusters in single copies in all plasmids (Cazares et al., 2020) were concatenated and used to generate a sequence alignment (MAFFT v7.490, BLOSUM 62; Geneious Prime v2023.0.1) from which a WAG substitution model (Jones et al., 1992) neighborjoining tree was generated. The bespoke R script associated with processing of the sequence data along with all output files are provided as Supplementary Material on figshare.

### Characterization of phenotypic and genomic concordance/discordance

For easier description and discussion of phenotypic and genomic results, we grouped the “susceptible, standard dosing regimen” (S) and “susceptible, increased exposure” (I) categories under the term “susceptible” as currently recommended by EUCAST. Whole-genome sequence (WGS) data were compared with DDT data for 31 *Pseudomonas* isolates against 10 antimicrobials (n = 310 combinations). For each combination, concordance was considered positive if a) WGS data were predicted to encode AMR genes and the isolate had a phenotypic resistant profile (WGS-R/DDT-R) or b) WGS data were not predicted to encode AMR genes and the isolate had a phenotypic susceptible profile (WGS-S/DDT-S) as described previously by (Rebelo et al., 2022; Vanstokstraeten et al., 2023). Discordance was considered positive in case of major or very major errors. Major errors (WGS-R/DDT-S) are defined as a resistant genotype and susceptible phenotype. Very major errors (WGS-S/DDT-R) are defined as a susceptible genotype and resistant phenotype. WGS results were classified as “resistant” when one or several AMR genes were identified by CARD and allocated as the mechanism of AMR to that antimicrobial, and as “susceptible” when no AMR gene was found.

## RESULTS

### Genome characterization

The draft genomes assembled from short-read and, in some instances, long-read data consisted of between 1 and 740 contigs. The average number of coding sequences predicted to be encoded within the genomes was 6,297 ± 289. Genomes had a mean G+C content of 66 %. The tRNA copy number for the isolates ranged from 59 to 70. All isolates were confirmed to be *P. aeruginosa* by ANI analysis against the genome of the type strain of *P. aeruginosa* (> 95–96 % ANI (Chun et al., 2018)), with additional support provided by phylogenetic analysis (**Supplementary Figure (1)**). The general features of the isolates’ genomes are provided in **Table (1)**.

### Genotypic and phenotypic AMR profiles

The AMR profiles of the 31 *P. aeruginosa* isolates were determined according to EUCAST guidelines. A summary of the classes of antimicrobials the isolates were resistant to is shown in **Figure (1)**. The isolates were highly resistant to quinolones [ciprofloxacin (n=12/31, 38.7 %) and levofloxacin (n=9/31, 29 %)] followed by tobramycin (n=10/31, 32.5 %) and cephalosporins (n=7/31, 22.5 %). Six (P5, P18, P20, P26, P28, P30) of the 31 isolates (19.3 %) were MDR (i.e. resistant to ≥ 3 antimicrobials from three different antibiotic classes) (**Table (3)**).

**Figure (1):**
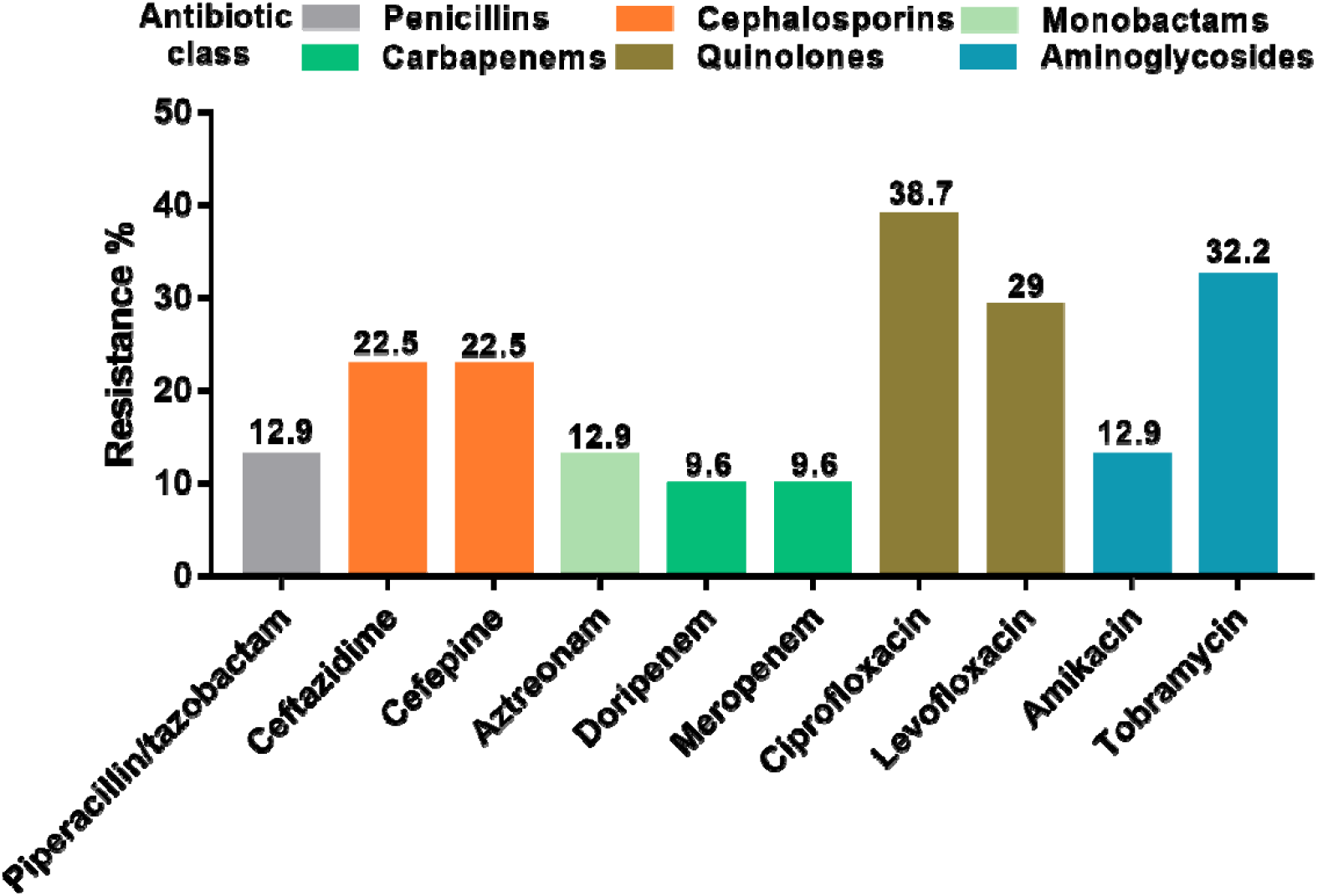
Classes of antimicrobials to which the 31 *P. aeruginosa* isolates recovered from CAUTIs were resistant. AMR susceptibility testing was done according to EUCAST guidelines. The figure depicts the proportion (%) of isolates that were resistant to each antibiotic.

**Table (3):** Overview for resistance genes of MDR isolates of *P. aeruginosa*. All isolates were predicted to encode the aminoglycoside-modifying enzyme *APH(3’)-IIb*.

| Isolate | β-lactamases | Resistance to fluoroquinolones | Others | Efflux pump systems | Phenotypic resistance profile * |
| --- | --- | --- | --- | --- | --- |
| P5 | *OXA-395*<br>*PDC-16* | *gyrA*<br>*qnrVC1* | *fosA*<br>*catB7*<br>*sul1* | MexAB-OprM<br>MexCD-OprJ<br>MexEF-OprN<br>MexHI-OpmD<br>MexPQ-OpmE | AK<br>ATM<br>CIP<br>FEP<br>LEV<br>TOB |
| P18 | *OXA-50*<br>*PDC-14* | – | *fosA*<br>*catB7* | MexAB-OprM<br>MexCD-OprJ<br>MexEF-OprN<br>MexHI-OpmD<br>MexPQ-OpmE | ATM<br>CIP<br>TOB<br>CAZ<br>FEP |
| P20 | *NDM-1*<br>*OXA-395*<br>*PDC-16* | *gyrA*<br>*qnrVC1* | *fosA*<br>*catB7*<br>*cmlA9*<br>*sul1*<br>*tet(D)* | MexAB-OprM<br>MexCD-OprJ<br>MexEF-OprN<br>MexHI-OpmD<br>MexPQ-OpmE | AK<br>CAZ<br>CIP<br>DOR<br>FEP<br>LEV<br>MEM<br>TOB<br>TZP |
| P26 | *NDM-1*<br>*OXA-395*<br>*PDC-16* | *gyrA*<br>*qnrVC1* | *fosA*<br>*catB7*<br>*cmlA9*<br>*sul1*<br>*tet(D)* | MexAB-OprM<br>MexCD-OprJ<br>MexEF-OprN<br>MexHI-OpmD<br>MexPQ-OpmE | AK<br>CAZ<br>CIP<br>DOR<br>FEP<br>LEV<br>MEM<br>TOB<br>TZP |
| P28 | *OXA-903*<br>*PDC-3* | – | *fosA*<br>*catB7* | MexAB-OprM<br>MexCD-OprJ<br>MexEF-OprN<br>MexHI-OpmD<br>MexPQ-OpmE | ATM<br>CAZ<br>CIP<br>FEP<br>TZP |
| P30 | *NDM-1*<br>*OXA-395*<br>*PDC-16* | *gyrA*<br>*qnrVC1* | *fosA*<br>*catB7*<br>*cmlA9*<br>*sul1*<br>*tet(D)* | MexAB-OprM<br>MexCD-OprJ<br>MexEF-OprN<br>MexHI-OpmD<br>MexPQ-OpmE | AK<br>CAZ<br>CIP<br>DOR<br>FEP<br>LEV<br>MEM<br>TOB<br>TZP |

\* AK, Amikacin; ATM, Aztreonam; CAZ, Ceftazidime; CIP, Ciprofloxacin; DOR, Doripenem; FEP, Cefepime; LEV, Levofloxacin; MEM, meropenem; TOB, Tobramycin; TZP, Piperacillin tazobactam.

Through genotypic analysis using RGI/CARD, a total of 88 antibiotic resistance genes were predicted to be encoded by the 31 isolates (726 perfect hits and 1182 strict hits), including genes conferring resistance to β-lactams, aminoglycosides, fluoroquinolones, macrolides and tetracyclines through different mechanisms, such as antibiotic efflux and antibiotic target alteration (*n* = 175), antibiotic inactivation (*n* = 179), antibiotic efflux (*n* = 1389), antibiotic target alteration (*n* = 80), reduced permeability to antibiotics (*n* = 62), antibiotic target protection (*n* = 10), and antibiotic target replacement (*n* = 13). RGI/CARD results for the *P. aeruginosa* isolates are summarized in **Figure (2)**, and compared with the phenotypic data.

**Figure (2):**
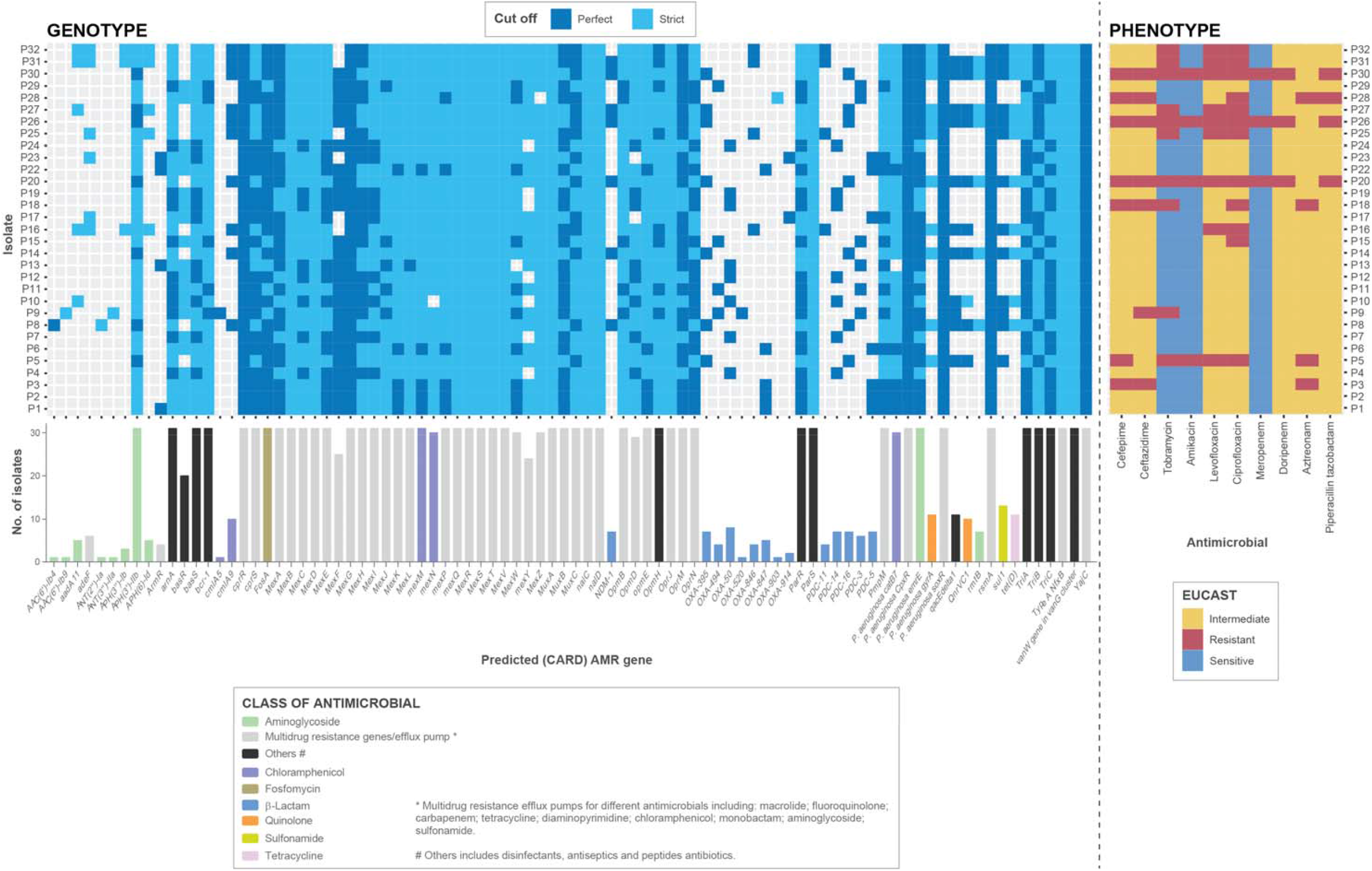
AMR genes predicted to be encoded within the genomes of the 31 isolates compared with their AMR phenotypic profiles (determined according to EUCAST guidelines). Resistomes were characterized using the RGI tool of CARD for perfect and strict hits. Strict CARD match, not identical but the bit score of the matched sequence is greater than the curated BLASTP bit score cut-off; perfect CARD match, 100_J % identical to the reference sequence along its entire length. The bar graphs under the genotypic data show the number of genomes encoding each predicted AMR gene. For the EUCAST data, intermediate refers to isolates considered “susceptible with increased exposure”.

In terms of comparing genotypic with phenotypic profiles for the MDR isolates, P5, P18, P20, P26, P28 and P30 were predicted to encode an aminoglycoside-modifying enzyme [APH(3’)-Ilb] and five efflux pump systems (MexAB-OprM, MexCD-OprJ, MexEF-OprN, MexHI-OpmD, and MexPQ-OpmE), while 4/6 and 5/6 of the MDR isolates were phenotypically resistant to the aminoglycosides amikacin and tobramycin, respectively. The genomes of isolates P20, P26 and P30 were also predicted to encode the β-lactamases *NDM-1, OXA-395*, and *PCD-16*; isolate P5 encoded *OXA-395* and PDC-16; isolate P18’s genome was predicted to encode *OXA-50* and *PDC-14*; isolate P28 was predicted to encode *OXA-903* and *PDC-3*. Phenotypically, 5/6 and 6/6 of the MDR isolates were resistant to ceftazidime and cefepime, respectively. Genes conferring resistance to quinolones (*gyrA* and *qnrVC1*) were predicted to be harbored by isolates P5, P20, P26 and P30 (**Table (3)**).

There were many additional resistance determinants predicted to be encoded within the genomes of the susceptible isolates with increased exposure (I): aminoglycoside-modifying enzymes *AAC(6’)-Ib4, AAC(6’)-Ib9, aadA11, ANT(2’’)-Ia, ANT(3’’)-IIa, APH(3’’)-Ib, APH(3’)-IIb, APH(6)-Id*; and the β-lactamases *OXA-50, OXA-395, OXA-494, OXA-520, OXA-846, OXA-847, OXA-903, OXA-914, PDC-3, PDC-5, PDC-11, PDC-14, PDC-16*” (**Figure (2)**).

Comparison of our WGS data and DDT results (with respect to predicted AMR genes and actual resistance phenotypes) yielded a concordance of 31 %, with discordant results (69 %) mainly due to phenotypically susceptible isolates predicted to encode AMR determinants in their genomes (e.g. isolate P29 concordant for resistance to pipracillin/tazobactam, but discordant for aztreonam; **Supplementary Table (1)**). However, the discordant cases were not even equally distributed. In 68.1 % of discordant cases, one or several AMR genes were predicted in the genome but the isolate was phenotypically susceptible (major errors, WGS-R/DDT-S; e.g. isolate P1 for the cephalosporins ceftazidime and cefepime). The remaining 0.9 % discordances were phenotypically resistant isolates in which no genetic determinants of AMR were predicted (very major errors, WGS-S/DDT-R; e.g. isolate P18 for the fluoroquinolone ciprofloxacin) (**Supplementary Table (1)**).

### Biofilm formation

Biofilm-forming abilities of the 31 isolates were tested, and compared with a known biofilm-negative control (*Salmonella enterica* serovar Enteritidis 27655S). *P. aeruginosa* isolates tended to form strong biofilms, with the isolates’ biofilm-forming ablilities classified as follows: non-biofilm producer (no change in OD_540_ over the medium control = 0.075), weak biofilm producer (up to a 2-fold change over the control), moderate biofilm producer (up to 4-fold change over the control), or strong biofilm producer (greater than 4-fold change over the control) (Stepanovic et al., 2000). The majority (77.4 %) of the isolates were strong biofilm-producers (P1, P3, P4, P5, P8, P9, P11, P12, P13, P14, P15, P17, P18, P19, P20, P22, P23, P25, P26, P27, P28, P30, P31, P32), 19.3 % were moderate (P2, P6, P7, P10, P16, P24), and 3.2 % were weak (P29) (**Figure (3)**).

**Figure (3):**
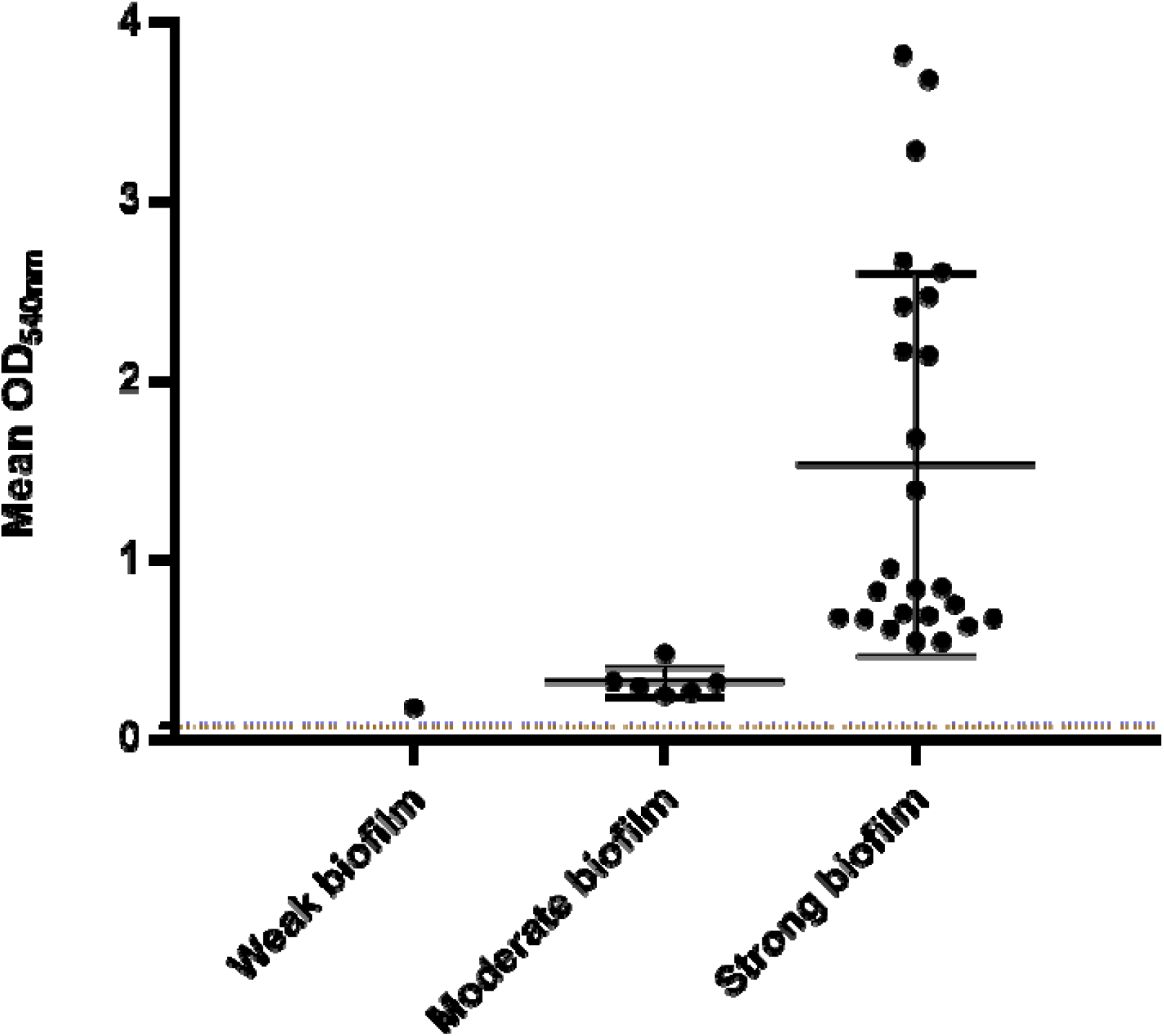
Classification of *P. aeruginosa* isolates according to their capacity to produce biofilm in TSBG. Data for each isolate are represented as the mean of four technical replicates (three biological replicates each). The blue dashed line (0.095) represents *Salmonella enterica serovar Enteritidis 27655S* while the brown dashed line (0.075) represents the medium. The mean and its standard deviation are represented for each biofilm formation category.

### Virulence factors associated with adherence and secretion systems

The investigation of virulence factors using VFDB predicted that isolates encode various virulence genes, ranging from 196 to 210 in number per isolate. Genes with no known functionality – “undetermined” in the VFDB database – were excluded from further analysis.

The major functional attributes of the known virulence factor genes detected in genomes were adherence (37.2 % abundance) and secretion systems (21.9 % abundance). All virulence genes detected by VFDB analysis are mentioned in **Supplmentary Table (2)**.

### MLST revealed multiple major clonal complexes

The clonal diversity among the 31 *P. aeruginosa* isolates showed eight different sequence types (STs): ST244, ST357, ST381, ST621, ST773, ST1430, ST1667 and ST3765 (**Table (1))**.

There were no relevant data in the PubMLST database regarding STs of *P. aeruginosa* in Egypt, although it is in the center of MENA region. We, therefore, compared the STs of the PubMLST database with those of our isolates, with respect to other countries and sources of infection (**Table (4)**). STs of *P. aeruginosa* in our study matched those of isolates detected outside the MENA region. PubMLST reported data for 107 ST244 isolates, 35 ST357 isolates, 47 ST381 isolates, four ST621 isolates, ten ST773 isolates, and one isolate each of ST1430, ST1667 and ST3765 across a range of non-MENA countries. Reported isolates of the MENA region had unique STs. The previous reported STs relevant to the MENA region are shown in **Table (5)**. The previous STs associated with UTIs are ST244 [Poland (4), Australia (1), Brazil (2)], ST357 [Poland (2)], and ST381 [Malaysia (1)].

**Table (4):** Summary of STs found in PubMLST database that matched those detected in this study. Bold text, associated with UTI.

| ST in current study | Source of isolation (*n* isolates) | Relevant countries (*n* isolates) |
| --- | --- | --- |
| ST244 | Blood (14)<br>Bronchial lavage (3)<br>Other (19)<br>Soft tissue infection (7)<br>Sputum (3)<br>**Urinary tract infection (7)**<br>Hospital effluent (3)<br>Water (2)<br>Soil (1) | Australia (10)<br>Brazil (12)<br>Central African Republic (3)<br>China (1)<br>France (10)<br>Ghana (1)<br>Ivory Coast (2)<br>Nigeria (2)<br>Poland (14)<br>Russia (3)<br>Spain (7)<br>UK (1)<br>Unknown (41) |
| ST357 | Bronchial lavage (6)<br>Water (1)<br>Other (5)<br>Soft tissue infection (2)<br>Sputum (2)<br>**Urinary tract infection (2)** | Brazil (2)<br>France (1)<br>Ghana (1)<br>Malaysia (2)<br>Nigeria (1)<br>Peru (4)<br>Poland (5)<br>Senegal (1)<br>Singapore (1)<br>Unknown (17) |
| ST381 | Blood (6)<br>Other (11)<br>Soft tissue infection (1)<br>Sputum (2)<br>Water (2)<br>Hospital effluent (1)<br>**Urinary tract infection (1)** | Australia (7)<br>Brazil (1)<br>France (4)<br>Ivory Coast (4)<br>Malaysia (2)<br>Poland (3)<br>Russia (3)<br>Spain (1)<br>Unknown (22) |
| ST621 | Unknown | Austria (1)<br>Unknown (3) |
| ST773 | Soft tissue infection (3)<br>Other (1)<br>Sputum (1)<br>Blood (1) | Bangladesh (1)<br>Central African Republic (1)<br>China (1)<br>Ghana (3)<br>Russia (1)<br>Unknown (3) |
| ST1430 | Unknown | Unknown (1) |
| ST1667 | Unknown | China (1) |
| ST3765 | Sputum (1) | Russia (1) |

**Table (5):** Summary for relevant STs found in PubMLST of *P. aeruginosa* in MENA region. Bold text, associated with UTI.

| Country | Source of infection (*n* isolates) | Relevant ST(s) |
| --- | --- | --- |
| Algeria | Blood (1) | 674 |
|  | Other (2) | 3349, 3350 |
| Iran | Soft tissue infection (2) | 967, 972 |
|  | Sputum (5) | 3118, 3119, 3377, 3381, 3382, 3450 |
|  | **Urinary tract infection (5)** | **970, 3376, 3378, 3379, 3380** |
| Iraq | Bronchial lavage (1) | 2209 |
|  | Other (2) | 2203, 2208 |
|  | Soft tissue infection (9) | 2196, 2197, 2198, 2199, 2200, 2201, 2202, 2205, 2206 |
|  | Sputum (2) | 2204, 2207 |
|  | **Urinary tract infection (3)** | **2195, 2210, 3352** |
| Kuwait | Unknown (1) | 3842 |
| Lebanon | Bronchial lavage (1) | 1702 |
|  | Other (5) | 1701, 1759, 1760, 1761, 1762 |
|  | **Urinary tract infection (3)** | **1699, 1700, 3425** |
|  | Unknown (1) | 3985 |
| Libya | Sputum (5) | 1924, 1925, 1926, 1927, 1928 |
| Palestine | Soft tissue infection (3) | 1562, 1563, 1564 |
| Saudi Arabia | Sputum (2) | 3728, 3729 |
|  | **Urinary tract infection (1)** | **3730** |
|  | Unknown (12) | 2010, 2012, 2013, 3710, 3711, 3712, 3713, 3714, 3715, 3716, 3717, 3718 |
| Sudan | Blood (2) | 3900 |
|  | **Urinary tract infection (3)** | **3898, 3899, 3901** |
| Tunisia | Other (11) | 2042, 2043, 2537, 2538, 3385, 3386, 3968, 3969, 3970 |
|  | Sputum (1) | 3762 |
|  | Water (1) | 2539 |
| Turkey | Blood (2) | 2529, 2531 |
|  | Bronchial lavage (1) | 2532 |
|  | Other (1) | 2034 |
|  | Soft tissue infection (15) | 2513, 2514, 2515, 2516, 2516, 2517, 2518, 2519, 2520, 2521, 2522, 2523, 2525, 2526, 2527 |
|  | **Urinary tract infection (2)** | **2528, 2530** |
| United Arab Emirates | Sputum (1) | 2011 |

### Megaplasmid identification

Visual inspection of Bandage maps (not shown) generated for our short-read draft genome assemblies suggested isolate P9 encoded a circular megaplasmid of >400,000 bp. The *repA, parA* and *virB4* sequences of megaplasmid pBT2436 were extracted from its sequence (accession CP039989) using the PCR primer sequences of (Cazares et al., 2020). These were used in a BLASTN search of the draft genomes for all our *P. aeruginosa* isolates. P9 alone returned hits, sharing 97.1 %, 99.4 % and 100 % similarity with the *repA, parA* and *virB4* nucleotide sequences, respectively. Confirmation of isolate P9 alone encoding a circular pBT2436-like megaplasmid was achieved by mapping the reads of all isolates against the genomes of the reference genomes (Cazares et al., 2020) listed in **Table (2)**. Between 10.01 % and 12.68 % of the Illumina reads of isolate P9 mapped to the pBT2436-like megaplasmid reference genomes (**Figure (5a)**). No other isolate had more than 1.8 % of their reads map to any of the reference megaplasmid sequences.

Consequently, a MinION/Illumina hybrid assembly was generated for P9 (**Table (1)**). The genome comprised a complete, circular chromosome (6,518,599 bp) and two complete, circular plasmids (pP9Me1, 422,938 bp; pP9Me2, 49,064 bp). The chromosome was predicted to encode 5,950 CDS. Neither plasmid matched sequences in PlasmidMLST. The megaplasmid pP9Me1 was assigned to PTU-Pse13 (score 1.000) by COPLA (Redondo-Salvo et al., 2021). pP9Me2 could not be assigned to a plasmid taxonomy unit using this tool. No mobility group, replication initiator protein domain or replicon type could be assigned to pP9Me1 or pP9Me2 by plaSquid. However, Bakta did identify a replication initiation protein (RepA) in pP9Me2’s sequence that shared homology with UniRef90_A0A218MAR0, a HK97 gp10 family phage protein of *P. aeruginosa*.

The megaplasmid pP9Me1 was predicted to encode 538 CDS, including the virulence genes (VFDB) *pilD* (type IV pili biosynthesis), *chpA* and *pilG* (type IV pili twitching motility-related proteins) and *csrA* (carbon storage regulator A), and the AMR genes *sul1, qacEdelta1*, OXA-520, *cmlA5* (CARD perfect matches) plus ANT(3’’)-Iia and AAC(6’)-Ib9 (CARD strict matches). Its sequence shared high similarity with that of pBT2436; a progressiveMauve alignment (not shown) of the sequences of pBT2436 and pM9Me1 showed them to share 163,628 identical sites (97 % pairwise identity), and they shared an average nucleotide identity (fastANI) of 98.5 % (**Figure (5b)**).

Plasmid pP9Me2 was predicted to encode 68 CDS; it did not encode any AMR-or virulence-associated genes based on CARD and VFDB searches. Based on an NCBI BLASTN analysis, its sequence shared high similarity with that of the circular and complete (50,754 bp; GenBank accession CP081288.1) *P. aeruginosa* plasmid pF092021-1 (93 % query coverage, 98.7 % identity; **Supplementary Figure (2)**). A progressive Mauve alignment of the sequences showed pP9Me2 and pF092021-1 to share 44,425 identical sites (81.1. % pairwise identity) (**Supplementary Figure (3)**); average nucleotide identity could not be determined for these plasmid sequences.

In their original study, (Cazares et al., 2020) identified 15 pBT2436-like megaplasmids (**Table (2)**). BLASTN searches (Supplementary Material: BLASTN_hits_plasmids.xlsx) of the pBT2436 *repA, parA* and *virB4* sequences against all complete *Pseudomonas* plasmid sequences >200,000 bp from NCBI Genome identified a further 24 potential pBT2436-like megaplasmids encoding only one copy each of the three pBT2436-like sequences (**Table (6)**). FastANI analysis showed the sequences of these plasmids shared between 95.9 and 100 % average nucleotide identity with one another, pP9Me1 and the 15 reference sequences (**Supplementary Figure (4)**). Consequently, the protein sequences predicted to be encoded by the 40 megaplasmids were clustered, to identify single-copy proteins that shared 80 % identity and 80 % coverage with the core sequences of pBT2436 (Cazares et al., 2020). Of the 261 core sequences described for pBT2436, 217 were included in our analysis. We found an alignment (55,243 aa) of these concatenated sequences to share between 97.4 % and 100 % identity, with the sequences of plasmids pWTJH12-KPC (CP064404) and pZPPH29-KPC (CP077978) identical to one another (they were from isolates recovered in the same hospital (Y. Li et al., 2022)). Phylogenetic analysis confirmed how closely related the newly identified megaplasmids were to pP9Me1 and the reference pBT2436-like plasmids (**Figure (6)**).

**Table (6):**
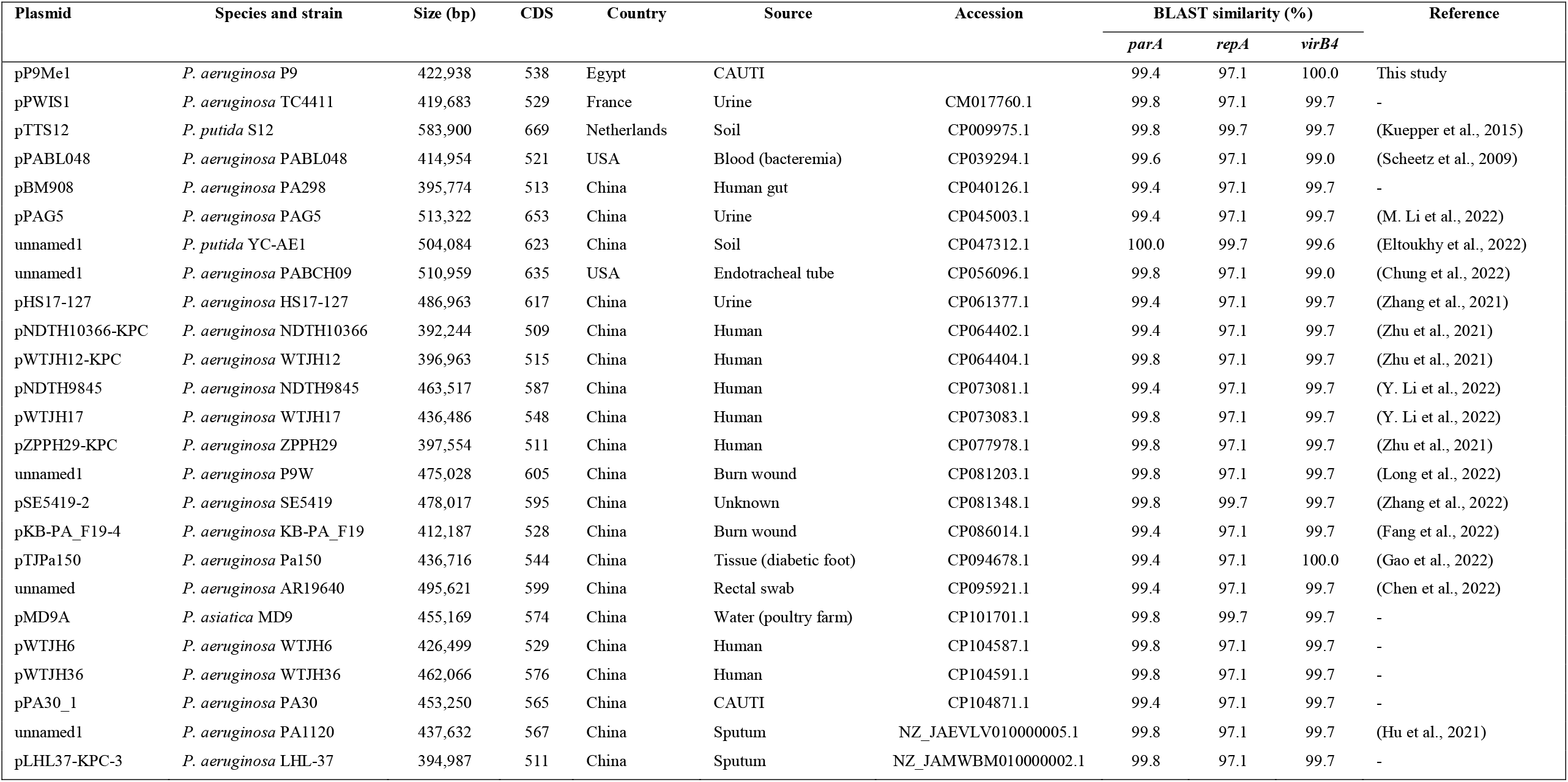
New *Pseudomonas* pBT2436-like megaplasmids identified in this study.

| Plasmid | Species and strain | Size (bp) | CDS | Country | Source | Accession | BLAST similarity (%) |  |  | Reference |
| --- | --- | --- | --- | --- | --- | --- | --- | --- | --- | --- |
|  |  |  |  |  |  |  | *parA* | *repA* | *virB4* |  |
| pP9Me1 | *P. aeruginosa* P9 | 422,938 | 538 | Egypt | CAUTI |  | 99.4 | 97.1 | 100.0 | This study |
| pPWIS1 | *P. aeruginosa* TC4411 | 419,683 | 529 | France | Urine | CM017760.1 | 99.8 | 97.1 | 99.7 | - |
| pTTS12 | *P. putida* S12 | 583,900 | 669 | Netherlands | Soil | CP009975.1 | 99.8 | 99.7 | 99.7 | (Kuepper et al., 2015) |
| pPABL048 | *P. aeruginosa* PABL048 | 414,954 | 521 | USA | Blood (bacteremia) | CP039294.1 | 99.6 | 97.1 | 99.0 | (Scheetz et al., 2009) |
| pBM908 | *P. aeruginosa* PA298 | 395,774 | 513 | China | Human gut | CP040126.1 | 99.4 | 97.1 | 99.7 | - |
| pPAG5 | *P. aeruginosa* PAG5 | 513,322 | 653 | China | Urine | CP045003.1 | 99.4 | 97.1 | 99.7 | (M. Li et al., 2022) |
| unnamed1 | *P. putida* YC-AE1 | 504,084 | 623 | China | Soil | CP047312.1 | 100.0 | 99.7 | 99.6 | (Eltoukhy et al., 2022) |
| unnamed1 | *P. aeruginosa* PABCH09 | 510,959 | 635 | USA | Endotracheal tube | CP056096.1 | 99.8 | 97.1 | 99.0 | (Chung et al., 2022) |
| pHS17-127 | *P. aeruginosa* HS17-127 | 486,963 | 617 | China | Urine | CP061377.1 | 99.4 | 97.1 | 99.7 | (Zhang et al., 2021) |
| pNDTH10366-KPC | *P. aeruginosa* NDTH10366 | 392,244 | 509 | China | Human | CP064402.1 | 99.4 | 97.1 | 99.7 | (Zhu et al., 2021) |
| pWTJH12-KPC | *P. aeruginosa* WTJH12 | 396,963 | 515 | China | Human | CP064404.1 | 99.8 | 97.1 | 99.7 | (Zhu et al., 2021) |
| pNDTH9845 | *P. aeruginosa* NDTH9845 | 463,517 | 587 | China | Human | CP073081.1 | 99.4 | 97.1 | 99.7 | (Y. Li et al., 2022) |
| pWTJH17 | *P. aeruginosa* WTJH17 | 436,486 | 548 | China | Human | CP073083.1 | 99.8 | 97.1 | 99.7 | (Y. Li et al., 2022) |
| pZPPH29-KPC | *P. aeruginosa* ZPPH29 | 397,554 | 511 | China | Human | CP077978.1 | 99.8 | 97.1 | 99.7 | (Zhu et al., 2021) |
| unnamed1 | *P. aeruginosa* P9W | 475,028 | 605 | China | Burn wound | CP081203.1 | 99.8 | 97.1 | 99.7 | (Long et al., 2022) |
| pSE5419-2 | *P. aeruginosa* SE5419 | 478,017 | 595 | China | Unknown | CP081348.1 | 99.8 | 99.7 | 99.7 | (Zhang et al., 2022) |
| pKB-PA_F19-4 | *P. aeruginosa* KB-PA_F19 | 412,187 | 528 | China | Burn wound | CP086014.1 | 99.4 | 97.1 | 99.7 | (Fang et al., 2022) |
| pTJPa150 | *P. aeruginosa* Pa150 | 436,716 | 544 | China | Tissue (diabetic foot) | CP094678.1 | 99.4 | 97.1 | 100.0 | (Gao et al., 2022) |
| unnamed | *P. aeruginosa* AR19640 | 495,621 | 599 | China | Rectal swab | CP095921.1 | 99.4 | 97.1 | 99.7 | (Chen et al., 2022) |
| pMD9A | *P. asiatica* MD9 | 455,169 | 574 | China | Water (poultry farm) | CP101701.1 | 99.8 | 99.7 | 99.7 | - |
| pWTJH6 | *P. aeruginosa* WTJH6 | 426,499 | 529 | China | Human | CP104587.1 | 99.8 | 97.1 | 99.7 | - |
| pWTJH36 | *P. aeruginosa* WTJH36 | 462,066 | 576 | China | Human | CP104591.1 | 99.8 | 97.1 | 99.7 | - |
| pPA30_1 | *P. aeruginosa* PA30 | 453,250 | 565 | China | CAUTI | CP104871.1 | 99.4 | 97.1 | 99.7 | - |
| unnamed1 | *P. aeruginosa* PA1120 | 437,632 | 567 | China | Sputum | NZ_JAEVLV010000005.1 | 99.8 | 97.1 | 99.7 | (Hu et al., 2021) |
| pLHL37-KPC-3 | *P. aeruginosa* LHL-37 | 394,987 | 511 | China | Sputum | NZ_JAMWBM010000002.1 | 99.8 | 97.1 | 99.7 | - |

## DISCUSSION

Genomes of *P. aeruginosa* are complex and highly variable, therefore various resistance genes can be acquired by them from non-fermentative bacteria or even from different strains of Enterobacterales. The genomic size ranges from 5.8 to 7.3 Mbp, with a core genome consisting of more than 4,000 genes plus a variable accessory gene pool (Arnold et al., 2015; Klockgether et al., 2011). *P. aeruginosa* is a tough bacterium to kill and it persists even after prolonged antibiotic treatment (Cottalorda et al., 2022; Cottalorda et al., 2021). It is recognized to encode an array of virulence factors and AMR genes that enable colonization and successful establishment of UTIs. In the MENA region, there is high-level resistance to antimicrobials in Iraq (100 %), Egypt (100 %), and Saudi Arabia (88.9 %) indicating difficulties in managing UTIs secondary to MDR *P. aeruginosa* (Al-Orphaly et al., 2021). However, prior to the current study, there were no data available on the genomic diversity of *P. aeruginosa* isolates associated with CAUTIs in Egypt. Through phenotypic and genotypic characterization of such isolates collected from an Egyptian hospital over a 3-month period, we have demonstrated MDR (**Table (3)**), high-risk clones of *P. aeruginosa* are present in this clinical setting. We have also identified the presence of a pBT2436-like megaplasmid in an Egyptian isolate of *P. aeruginosa*.

*P. aeruginosa* high-risk clones are disseminated worldwide and are common causative agents of HAIs. A common feature of high-risk clones is their ability to express β-lactamases and metallo-β-lactamases. The emergence of MDR *P. aeruginosa* is considered a significant public health issue (Angeletti et al., 2018). MDR, internationally important *P. aeruginosa* high-risk clones include ST111, ST175, ST233, ST235, ST277, ST357, ST654, and ST773 (Kocsis et al., 2021). We identified eight different STs among the CAUTI isolates characterized in this study, including the high-risk clones ST357 (n=4) and ST773 (n=7), neither of which has been reported previously in Egypt (**Table (4)**). The only previous reported ST in tertiary care Egyptian hospitals for *Pseudomonas* was ST233 (wound, sputum, urine and ear-swab samples), found to encode *NDM-1* and/or *VIM-2* by PCR (Zafer et al., 2015; Zafer et al., 2014). Our ST357 isolates (P16, P25, P31 and P32) were predicted to encode perfect sequence matches to the class C and D β-lactamases *PDC-11* and *OXA-846*, respectively. None was MDR based on phenotypic analysis, but they all showed susceptibility with increased exposure to the β-lactams [i.e. penicillin (piperacillin -tazobactam), cephalosporins (cefepime, ceftazidime), monobactam (aztreonam) and carbapenems (doripenem, meropenem)] tested (**Figure (2)**). The seven ST773 isolates (P5, P8, P14, P20, P26, P27 and P30) were all predicted to encode perfect matches to *PDC-16* and *OXA-395*, with all except P5 also encoding a perfect match to the metallo-β-lactamase *NDM-1*; isolates P5, P20, P26 and P30 were considered MDR based on EUCAST testing (**Figure (2), Table (3)**).

While PubMLST did not report data for ST357 in the MENA region (**Table (5)**), this sequence type has been reported in Qatar (bloodstream infections, clinical isolates), Lebanon (clinical infections), Bahrain (clinical isolates) and Saudi Arabia (bacteremia, clinical isolates) (Alamri et al., 2020; Bitar et al., 2022; Sid Ahmed et al., 2022; Sid Ahmed et al., 2020; Zowawi et al., 2018). ST773 has only previously been reported as a clone disseminated in a burns unit in Iran (Yousefi et al., 2013). Based on data available from PubMLST, ST357 has only once before been associated with UTIs (**Table (4)**), while this study is the first to report ST773 associated with a CAUTI. Our ST data will be deposited in the PubMLST database to add to information available from the MENA region and to facilitate tracking of clinically important *P. aeruginosa* isolates contributing to infections (**Table (5)**).

Many factors are responsible for the inherent antimicrobial resistance of *P. aeruginosa*: a large and adaptable genome, mobile genetic elements, a cell wall with low permeability and the ability of the bacterium to form biofilms (Lambert, 2002). Megaplasmids (plasmids >350 kbp in *Pseudomonas* (Hall et al., 2022)) are of emerging interest in the context of clinical infections associated with *P. aeruginosa*, as they have been found in nosocomial populations, are often self-transmitting and can encode a range of virulence and AMR genes (Urbanowicz et al., 2021). Plasmid pBT2436, although >420 kbp in size, can transmit multiple resistance determinants at high efficiency (Cazares et al., 2020). We identified a pBT2436-like megaplasmid (pP9Me1, 422,938 bp) within the genome of isolate P9 (ST3765). None of the other ST3765 isolates (P11, P15, P29) we characterized harbored pBT2436-like megaplasmids nor did any of our other isolates based on BLASTN and read-mapping analyses (**Figure 5(a)**). pP9Me1 encoded a range of virulence factors (*pilD, chpA, pilG, csrA*). Isolate P9 was determined to be a strong biofilm-former by phenotypic analysis; whether virulence genes encoded by pP9Me1 contribute to this phenotype will be the subject of future work. Similar to other pBT2436-like megaplasmids (Cazares et al., 2020), pP9Me1 encoded a range of AMR genes; the most notable of these was OXA-520, which belongs to the OXA-10 family of class D β-lactamases and has not been reported in Egypt previously. While included in the CARD RGI database we have been unable to find *Pseudomonas* reports on OXA-520 in Egypt, but it has reported in the Netherlands (Croughs et al., 2018; del Barrio-Tofiño et al., 2020).

Along with the megaplasmid pP9Me1, we identified a novel plasmid (pP9Me2, 49,064 bp) within the genome of isolate P9. This smaller plasmid is predicted to encode several putative conjugation genes. Whether pP9Me1 is transmissible and pP9Me2 contributes to this transmissibility will be the subject of future studies.

Complete *Pseudomonas* plasmid sequences deposited with NCBI Genome were searched for genes homologous to core protein sequences from pBT2436 using a combination of BLASTN-based (**Table (6)**), average nucleotide (**Supplementary Figure (4)**), and phylogenetic analyses (**Figure (6)**). We identified an additional 24 pBT2436-like megaplasmids and have extended the range over which they have been found: in addition to these plasmids having been detected in Thailand, China, Portugal, Switzerland (Cazares et al., 2020) and Egypt (this study), they can be found in the USA (n=2), Netherlands (n=1) and France (n=1) (**Table (6)**). To date, pBT2436-like megaplasmids have been detected in urine (n=3), CAUTIs (n=2) and UTIs (n=1) in China, France and Egypt (**Table (2), Table (6)**).

Efflux pumps are of great concern with respect to the emergence of antibacterial resistance in *P. aeruginosa* (Blanco et al., 2016; Kishk et al., 2020). Empirical therapy refers to the initiation of treatment before the results of diagnostic tests (such as bacterial culture and susceptibility testing) are available. When it comes to UTIs caused by *Pseudomonas* spp., empirical therapy can be challenging because of the potential for multidrug-resistance among these bacteria. In Egypt, empirical therapy for UTIs typically includes the use of fluoroquinolones (ciprofloxacin and levofloxacin) (Abdelkhalik et al., 2018; Nouh et al., 2021). These antibiotics are broad-spectrum and have good activity against *Pseudomonas*, although nearly 40 % of isolates in our study were resistant to ciprofloxacin. Other antibiotics such as cephalosporins (ceftazidime) and aminoglycosides (tobramycin) also can be used (Moustafa et al., 2021). It is also important to note that empirical therapy should only be used as a temporary measure, and that definitive therapy should be based on the results of bacterial culture and susceptibility testing. The choice of antimicrobial therapy should be guided by spectrum and susceptibility patterns of the etiological pathogens, tolerability and adverse reactions, costs, and availability.

Our study showed 22.5 % resistance to cephalosporins among the 31 isolates characterized, but a higher resistance was observed with quinolones (**Figure (1)**). This high resistance associated with quinolones is due to antibiotic misuse by patients as they are easily bought without prescription in Egypt (Ramadan et al., 2019). Comparing the antimicrobial susceptibility seen in this study with that in other countries in the MENA region, ciprofloxacin demonstrated high resistance in Bahrain (100 %), Tunisia (100 %), Qatar (91.2 %), Libya (91 %), Egypt (70 %), Jordan (50.9 %), Yemen (35.7 %), Lebanon (27 %), Iraq (22.7 %), Saudia Arabia (18.1 %), and Oman (15 %). The 3^rd^ and 4^th^ generation antipseudomonal cephalosporins demonstrated exceptionally high resistance within MDR *P. aeruginosa* clinical isolates in Qatar (96.6 %), Bahrain (86 %), Tunisia (70 %), Egypt (68 %), Libya (66 %), Yemen (47.1 %), and Iraq (41.2 %) (Al-Orphaly et al., 2021).

Susceptibility with increased exposure was seen for 90 % (doripenem) and 87 % (piperacillin-tazobactam and aztreonam) of our isolates (**Supplementary Table (1)**). The “I” susceptibility category was devised so patients infected by intermediate susceptible bacteria would be treated with a high dose of the relevant drug (Rodloff et al., 2008). MexAB-OprM is a multidrug efflux protein expressed in *P. aeruginosa*. MexA is the membrane fusion protein, MexB is the inner membrane transporter, and OprM is the outer membrane channel (Tsutsumi et al., 2019). Four active efflux pumps may be responsible for an increased (2-to 16-fold) resistance to fluoroquinolones when overexpressed; namely, MexAB-OprM, MexXY/OprM, MexCD-OprJ, and MexEF-OprN (Köhler et al., 1997; Masuda et al., 2000; Zhang et al., 2001).

Other efflux systems MexHI-OpmD and MexPQ-OpmE have also been reported to export fluoroquinolones in *P. aeruginosa* (Mima et al., 2005; Sekiya et al., 2003). In our study, as shown in **Figure (2)**, all isolates harbored multiple genes responsible for the mentioned efflux-pump systems. Overexpression of efflux pumps could be the leading cause of MDR in bacteria as it leads to a decreased intracellular concentration of antibiotics and reduced susceptibility to antimicrobial agents due to continuous expelling of structurally unrelated drugs (Khosravi & Mihani, 2008).

Genotypic detection of resistance determinants revealed that all isolates were predicted to encode numerous AMR genes (**Figure (2)**) associated with resistance to aminoglycosides [*AAC(6’)-Ib4, AAC(6’)-Ib9, aadA11, aadA2, ANT(2’’)-Ia, ANT(3’’)-IIa, APH(3’)-Ia, APH(3’’)-Ib, APH(3’)-IIb, APH(6)-Id*], β-lactamases (*NDM-1, PDC-3, PDC-5, PDC-11, PDC-14, PDC-16, OXA-50, OXA-395, OXA-494, OXA-520, OXA-846, OXA-847, OXA-903, OXA-914*), fluoroquinolones (*gyrA, qnrVC1*), fosfomycin (*fosA*), sulfonamides (*sul1, sul2*), tetracyclines [*tet(C), tet(D)*] and chloramphenicol (*cmlA5, cmlA9, mexM, mexN, catB7*). However, resistance determinants mentioned in previous Egyptian reports, namely *AmpC, IMP* and *VIM* (Abbas et al., 2018; Basha et al., 2020; El-Domany et al., 2017), were not detected in the current study.

While the β-lactamases *OXA-2, OXA-4, OXA-10, OXA-50, OXA-486* and *PDC-3* have been reported for *P. aeruginosa* from urine, intensive care unit-associated infections, and general infections in Egypt, Saudia Arabia and Qatar (Al-Agamy et al., 2016; El-Shouny et al., 2018; Sid Ahmed et al., 2020), the current study is the first to report the presence of *OXA-395, OXA-494, OXA-520* (discussed above), *OXA-846, OXA-847, OXA-903, OXA-914, PDC-5, PDC-11, PDC-14*, and *PDC-16* in *P. aeruginosa* in Egypt.

There are discrepancies in the literature when comparing genomic and phenotypic data for *Pseudomonas* spp. and other bacteria contributing to infections. In a recent study, the highest discordance between predicted AMR genes and phenotypic resistance profiles was observed with *P. aeruginosa* isolates (n =21; 9 antimicrobials, 189 combinations) rather than other Enterobacterales or Gram-positive bacteria (Rebelo et al., 2022); 44.4 % of the results for the *P. aeruginosa* isolates showed discordance between phenotype and genotype. A third (63/189) of discordant results were major errors and 11.1 % (21/189) were very major errors. Worth mentioning is that 11 of the *P. aeruginosa* isolates showing discordant results were isolated from urine (Rebelo et al., 2022). Another recent study showed that isolates recovered from urine produced greatest discordance between genomic and phenotypic data for AMR profiles of both Enterobacterales and *P. aeruginosa*. Clinical implications could be drastic if hospitals are relying on “*susceptibility of one carbapenem to confer susceptibility to another carbapenem*” when interpreting data (Ku et al., 2021).

We suggest our high discordance level (i.e. major errors WGS-R/DDT-S; 68.1 %) may be accounted for due to pooling of “S” and “I” isolates together into one category in accordance with the EUCAST update for susceptibility definitions in 2019. Because of these new definitions and breakpoints, *P. aeruginosa* becomes intrinsically less susceptible to an antimicrobial, and will thus never reach the “S” susceptible category. Infections require increased exposure for almost all antimicrobials to be treated, hence *P. aeruginosa* phenotypes fall into the clinical category of “susceptible with increased exposure” (i.e. “I”) for all relevant antimicrobials (except meropenem) (Nabal Díaz et al., 2022). An in-depth review of genotype-phenotype AMR concordance was done by the EUCAST subcommittee, which concluded that promising high levels of concordance were noted for certain bacterial groups (*Enterobacteriaceae* and staphylococci), while other species (*P. aeruginosa* and *Acinetobacter baumannii*) proved much more difficult to interpret (Ellington et al., 2017). The major challenge for *P. aeruginosa* and *A. baumannii* lies in the identification or prediction of resistance due to chromosomal alterations resulting in modification of expression levels, particularly with respect to efflux pumps, outer membrane proteins and intrinsic β-lactamases.

For many bacteria, the urinary tract represents a harsh, nutrient-limited environment; thus, to survive and grow within the urinary tract, *P. aeruginosa* produces toxins and proteases that injure the host tissue to release nutrients, while also providing a niche for bacterial invasion and dissemination (Flores-Mireles et al., 2015). As shown in **Figure (4)** and mentioned in **Supplementary Table (2)**, our isolates encoded genes predicted to produce proteases, toxins, quorum sensing and secretion systems. The main traits of the virulence genes predicted to be encoded by the isolates characterized in this study were related to adherence and secretion systems, thus signifying that the isolates could be biofilm-producers as suggested by a previous report (Datar et al., 2021). The process of biofilm formation in *P. aeruginosa* is complex and multifactorial, involving the coordination of many different genes including those encoding for motility, quorum sensing, alginate production and regulation systems (Redfern et al., 2021; Thi et al., 2020).

**Figure (4):**
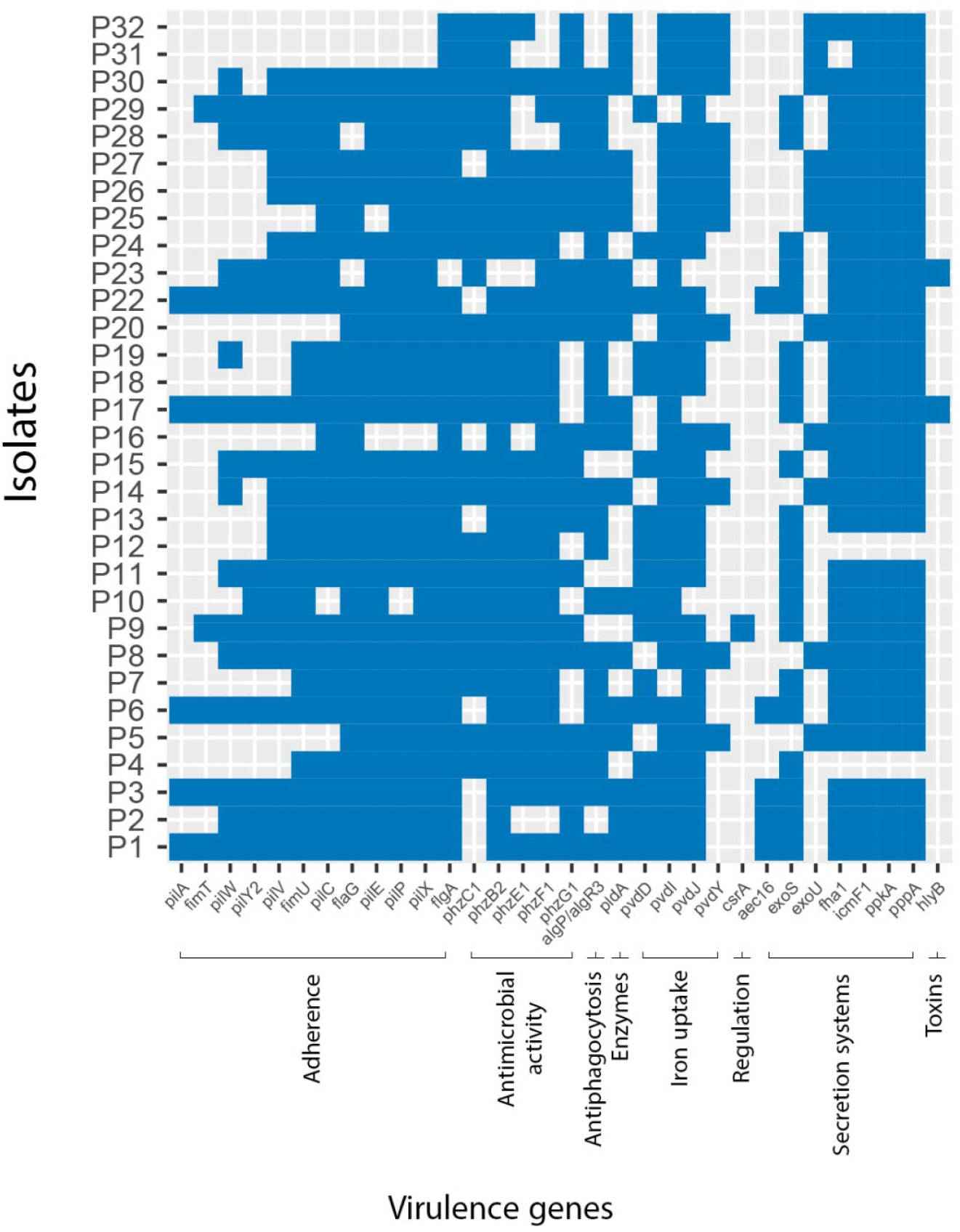
Prevalence of virulence factors (<100 % presence) predicted to be encoded within the genomes of the 31 *P. aeruginosa* isolates using the VFDB. Adherence: *pilA, fimT, pilY2, pilW, pilV, fimU, pilC, flaG, pilE, pilP, pilX, flgA*. Antimicrobial activity: *phzC1, phzG1F1, phzB2*. Antiphagocytosis: *algP/algR3*. Enzymes: *pldA*. Iron uptake: *pvdY, pvdD, pvdJ, pvdI*. Regulation: *csrA*. Secretion systems: *aec16, exoU, exoS, fha1, icmF1, ppkA, pppA*. Toxins: *hlyB*.

In comparison with a previous report (Díaz-Ríos et al., 2021), a total of 220 virulence genes were found among their *Pseudomonas* biofilm-forming isolates by comparing their WGS and VFDB data. All the isolates were able to produce biofilm. The most-represented groups of virulence genes identified among the isolates’ genomes were those for flagellar protein synthesis (17 %), type III secretion system (T3SS) machinery (17.7 %), type IV pili-related functions and twitching motility (14.5 %), and alginate biosynthesis and regulation (12 %). In our study, a total of 215 of virulence genes [**Supplementary Table (2)**] were found, with most of our isolates forming a strong biofilm (**Figure (3)**). The most represented groups of virulence genes identified were those associated with flagellar protein synthesis (22.2 %), T3SS (18.5 %), type IV pili and twitching motility (14.8 %), and alginate biosynthesis and regulation (12.1 %).

*pilA* and *fimT* have previously been reported as biofilm-associated genes (Deligianni et al., 2010; Sultan et al., 2021). Another report showed MDR biofilm-forming *P. aeruginosa* ST111 encoded both *pilA* and *fimT*, but these genes were absent from the ST235 pan-genome. In our study, *pilA* and *fimT* genes were predicted to be encoded in the genomes of the strong biofilm-formers (P1, P3, P17, P22) and one of the moderate biofilm-formers (P6). *fimT* gene was found without *pilA* in isolates P9 and P29, which were strong and weak biofilm-formers, respectively. T3SS genes *exoT* and *exoY* were found in all isolates whereas *exoS* and *exoU*, were not found concurrently in our isolates; *exoU*^+^ isolates were P5, P8, P14, P16, P20, P25, P26, P27, P30, P31 and P32, while *exoS*^+^ isolates were P1, P2, P3, P4, P6, P7, P9, P10, P11, P12, P13, P15, P17, P18, P19, P22, P23, P24, P28, P29 (**Figure (5)**). In general, *Pseudomonas* encoding *exoS* and *exoT* show an invasive phenotype while those isolates encoding *exoU*, are cytotoxic in nature (Karthikeyan et al., 2013). *exoS* and e*xoU* are generally mutually exclusive, although some studies have reported rare isolates harboring both exotoxins (Rodrigues et al., 2020; Sarges et al., 2020).

**Figure (5):**
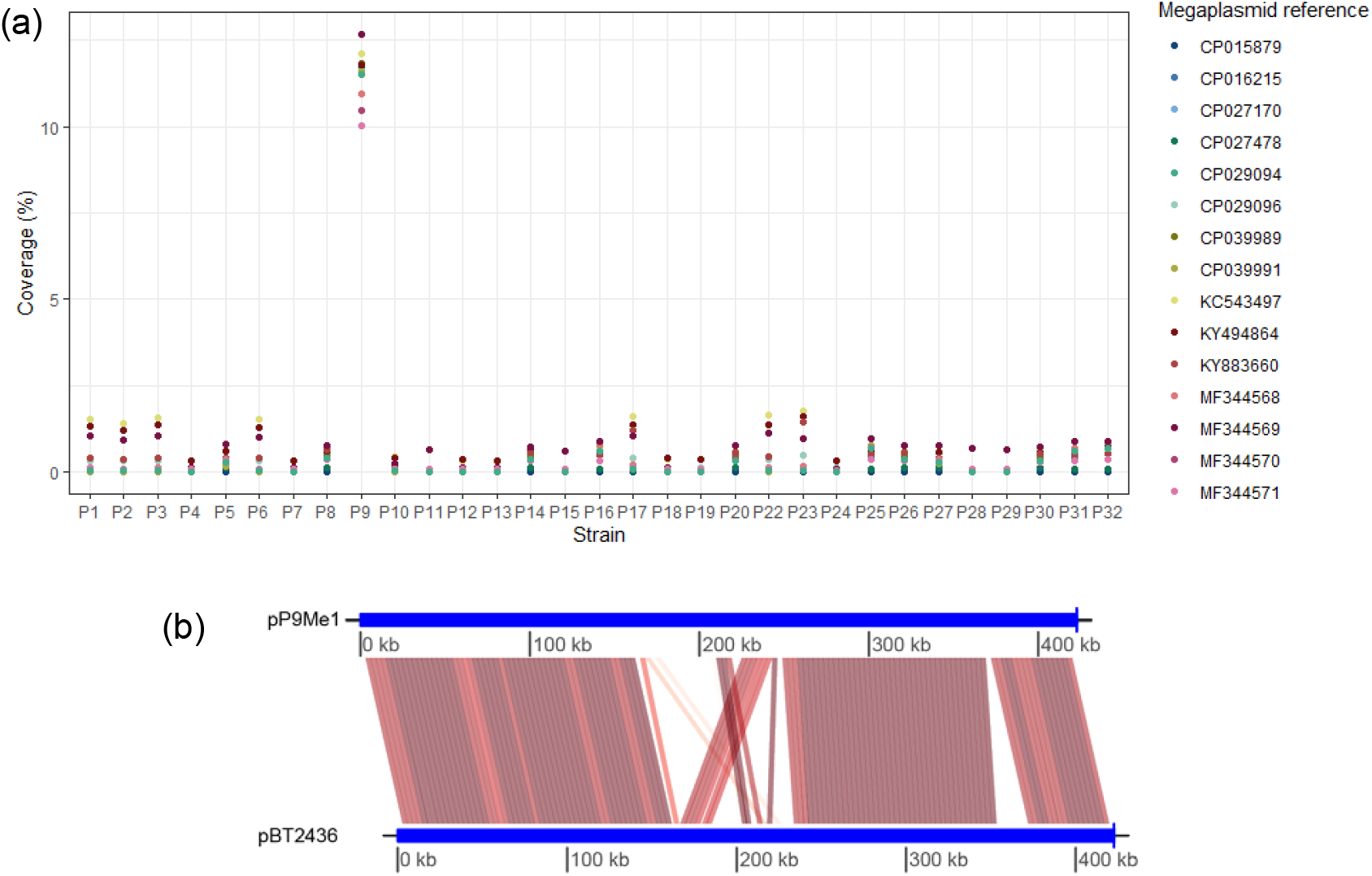
Detection and characterization of a pBT2436-like megaplasmid in the genome of *P. aeruginosa* P9. (a) Proportion of Illumina sequence reads generated for *P. aeruginosa* isolates recovered in Egypt that map to pBT2436-like megaplasmid reference sequences. (b) Visualization of the conserved regions between the sequences of the megaplasmids pP9Me1 and pBT2436 as determined using FastANI, with *repA* set as the start gene for both plasmid sequences.

**Figure (6):**
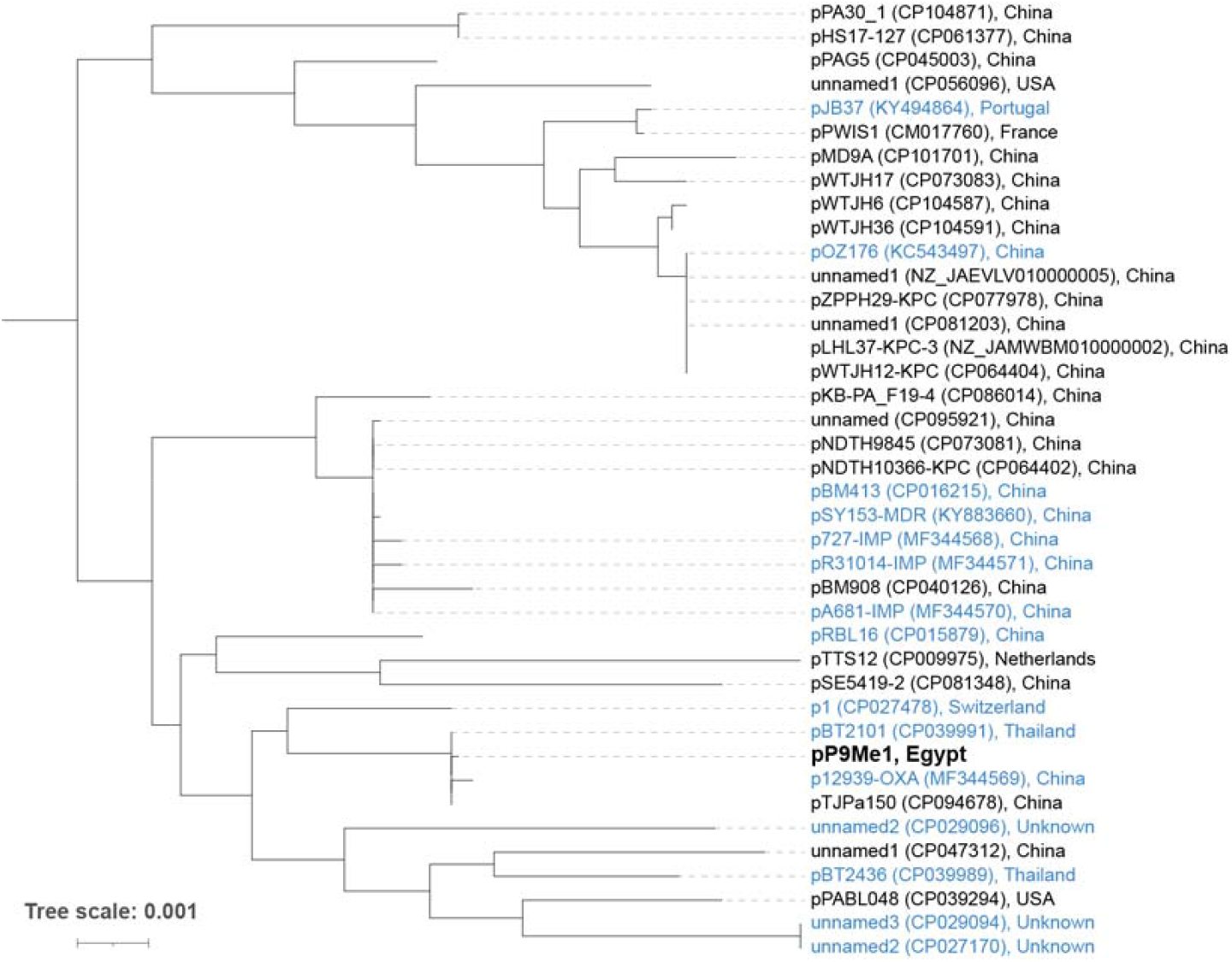
Phylogenetic (neighbour joining) tree showing relationships of megaplasmid pP9Me1 and other pBT2436-like megaplasmids. The tree, rooted at the midpoint, was built from a multiple-sequence alignment of 55,243 aa, comprising the sequences of 217/261 core proteins described by (Cazares et al., 2020). Plasmids shown in blue were defined as pBT2436-like by (Cazares et al., 2020), while those in black were identified as pBT2436-like in the current study. Scale bar, average number of amino acid substitutions per position. The three-clade structure for *Pseudomonas* plasmids seen here is similar to that generated by (Ramazanzadeh, 2021) for an analysis of 15 complete plasmid sequences of *P. aeruginosa*.

## CONCLUSIONS

This study demonstrates the utility of next-generation sequencing to define the diversity of AMR and virulence elements and highlight STs of *P. aeruginosa* contributing to CAUTIs in Egypt. This information is valuable in furthering the design of diagnostics and therapeutics for the treatment of *P. aeruginosa* infections in the MENA region. Continuous monitoring and surveillance programs should be encouraged in Egypt to track new high-risk clones and to analyze emergence of new clones as well as novel resistance determinants.

## Abbreviations

AMR: antimicrobial resistance
ANI: average nucleotide identity
CARD: Comprehensive Antibiotic Resistance Database
CAUTI: catheter-associated urinary tract infection
DDT: disc diffusion test
HAI: healthcare-associated infection
MDR: multidrug-resistant
MENA: Middle East and North Africa
RGI: Resistance Gene Identifier
ST: sequence type
T3SS: type 3 secretion system
TSBG: tryptone soy broth supplemented with glucose
UTI: urinary tract infection
VFDB: Virulence Factor Database
WGS: whole-genome sequence.

## DATA AVAILABILITY STATEMENT

Genome sequences for all samples used in this study have been deposited in National Centre for Biotechnology Information (NCBI) and are available under BioProject ID PRJNA913392.

## ACKNOWLEDGEMENTS

This work was supported by The Egyptian Ministry of Higher Education & Scientific Research represented by The Egyptian Bureau for Cultural & Educational Affairs in London. We would like to thank the Urology and Nephrology Centre, Mansoura, Egypt for providing the clinical isolates used in this study. We thank the Animal and Plant Health Agency, Addlestone, Surrey, UK for providing *Salmonella enterica* serovar Enteritidis 27655S to us under a Material Transfer Agreement. We thank Dr Gareth McVicker for providing guidance on the analysis of megaplasmid sequences.

ME – did all phenotypic work; extracted DNA for sequencing; characterized the AMR and virulence genes encoded by the isolates and their plasmids; MLST analysis and summary; interpreted virulence and AMR data. JCT – MinION sequencing and hybrid genome assembly. LH – annotated all genomes; did all phylogenetic analyses and megaplasmid bioinformatics; supervised the study. All authors contributed to writing of the manuscript and approved the final version.

## Notes

https://figshare.com/projects/Phenotypic_and_genomic_characterization_of_Pseudomonas_aeruginosa_isolates_recovered_from_catheter-associated_urinary_tract_infections_in_an_Egyptian_hospital/156639

### Competing Interest Statement

The authors have declared no competing interest.

