## Supplementary Figures for "Phenotypic and genomic characterization of *Pseudomonas aeruginosa* isolates recovered from catheter-associated urinary tract infections in an Egyptian hospital"

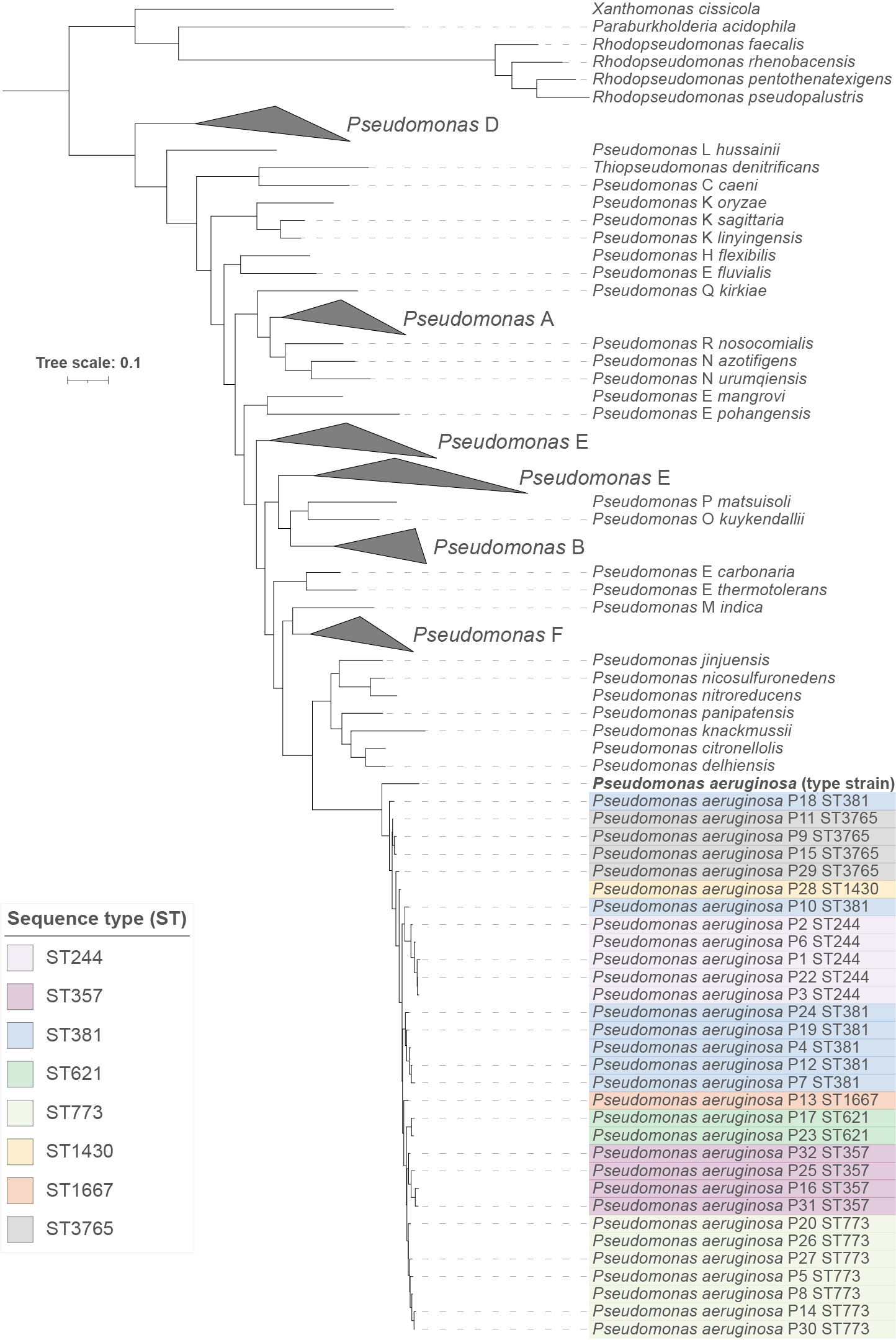

**Supplementary Figure (1):** Phylogenetic tree confirming the affiliation of the 31 clinical isolates from Egypt with *Pseudomonas aeruginosa*. Using PhyloPhlAn 3.0, the tree was generated from the proteomes of the 31 isolates and those of 245 reference sequences (GTDB species representatives, NCBI type material) downloaded from the Genome Taxonomy Database. The tree was annotated using iTOL v6 (Letunic & Bork, 2021) and Adobe Illustrator. The clinical isolates are colored according to the ST they belong to. Scale bar, average number of amino acid substitutions per position.

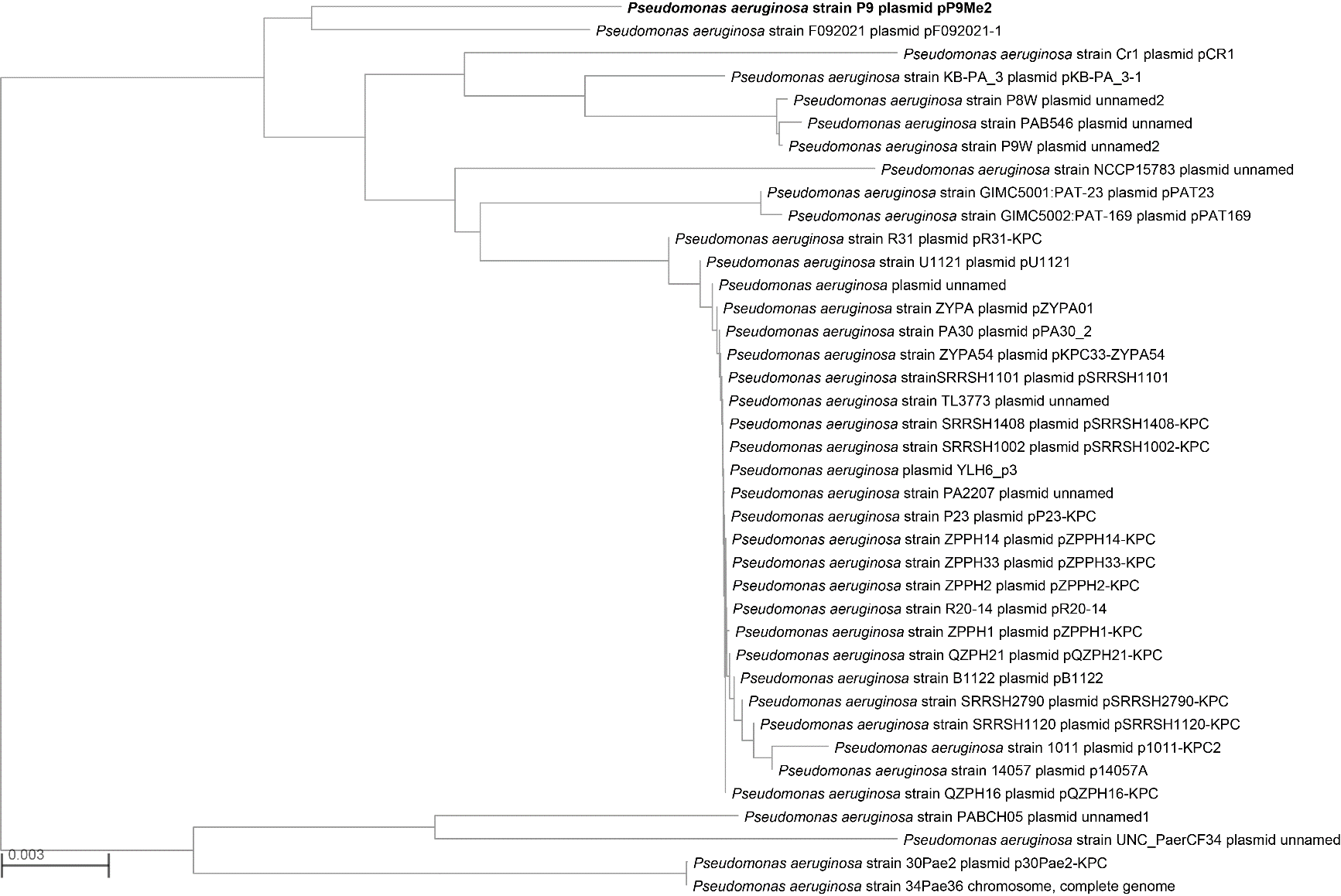

**Supplementary Figure (2):** Distance (neighbor-joining) tree generated by NCBI from BLASTN analysis of the sequence of p9Me2 with its closest relatives. The tree was rooted at the midpoint.

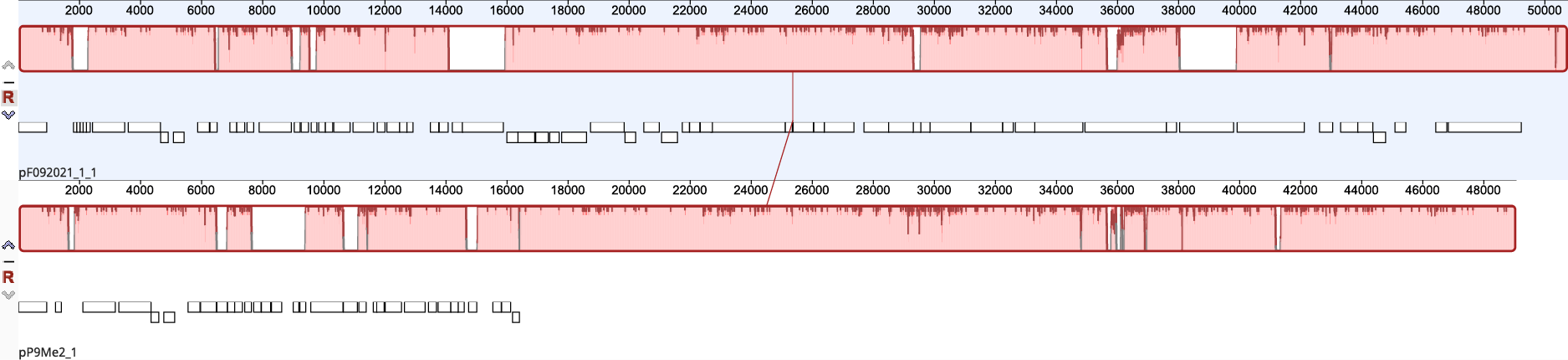

**Supplementary Figure (3):** progressiveMauve alignment of the sequences of plasmids pP9Me2 and pF09202-1, created using Geneious Prime v2023.0.1.

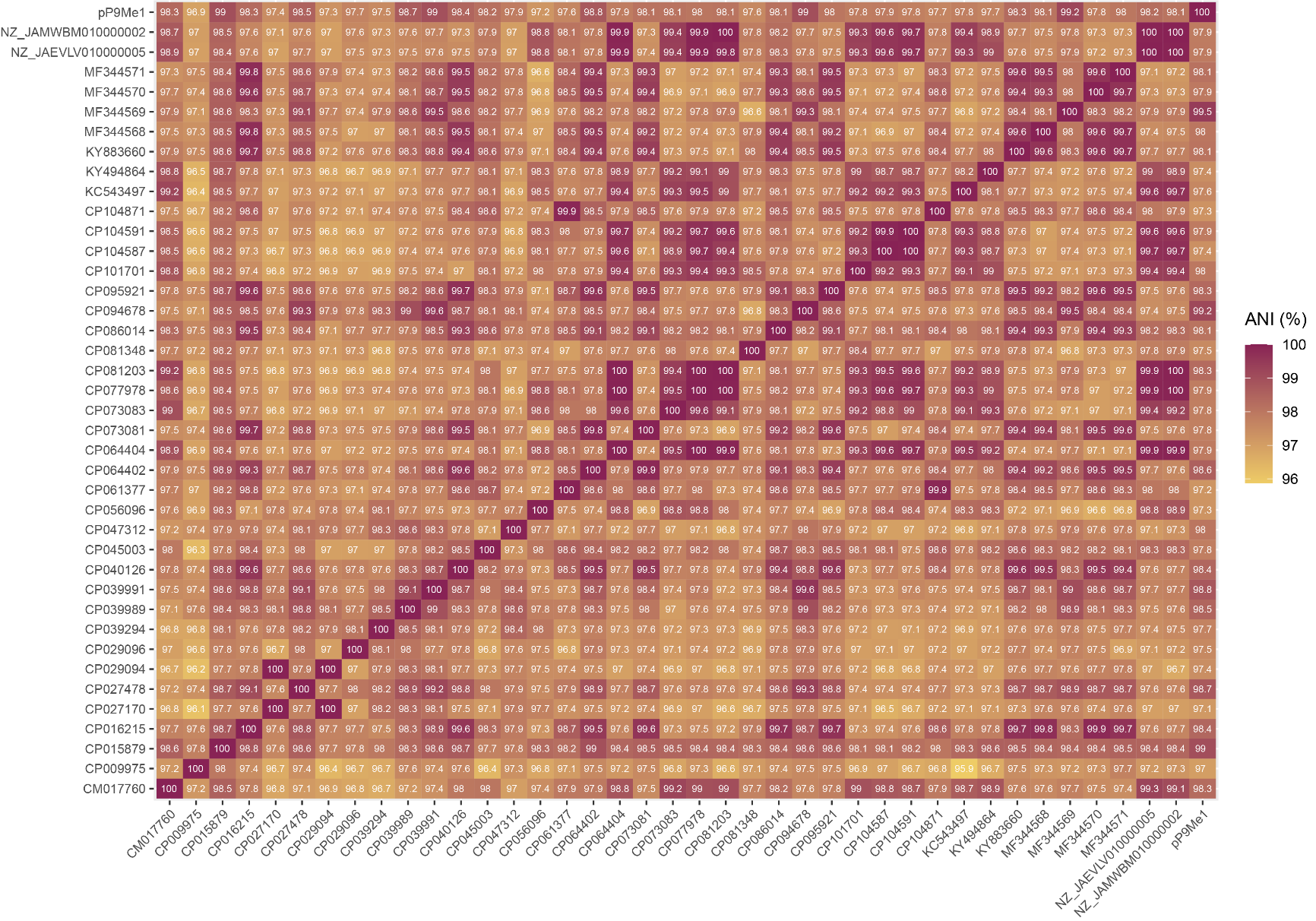

**Supplementary Figure (4):** Visualization of all-against-all FastANI analysis of the complete sequences of 40 pBT2436-like megaplasmids. Full details for the plasmids included in this analysis can be found in **Table (2)** and **Table (7)**.
